# On the state of chromatin in mammalian sperm

**DOI:** 10.64898/2026.07.09.737589

**Authors:** Katerina Tsimaratou, Victor G. Corces

**Affiliations:** Department of Human Genetics, Emory University School of Medicine, Atlanta, GA 30322

**Keywords:** Transcription, CTCF, Cohesin, Fertilization, Development

## Abstract

Mammalian sperm chromatin carries epigenetic information with the potential to influence offspring phenotype, making its faithful characterization essential. It has been suggested that cauda sperm preparations are contaminated by somatic chromatin, that this contamination dominates genome-wide profiles, and that valid results require pretreatment with somatic cell lysis buffer, DNase I, and dithiothreitol. Here we show that properly purified cauda sperm contain no detectable somatic cells or cell-free DNA and that this pretreatment disrupts sperm chromatin organization. SCLB permeabilizes the sperm nucleus, allowing DNase I to fragment the sperm genome in situ, while DTT treatment causes chromatin to leak out of the nucleus. Using ATAC-see, we further demonstrate that Tn5 transposase can access intact protamine-condensed sperm chromatin without DTT, refuting the premise that profiles from untreated sperm reflect contamination. Pretreatment therefore damages the chromatin it claims to purify, and published profiles of untreated cauda sperm are valid and require no systematic re-examination.

**HIGHLIGHTS:** Properly purified cauda sperm show no detectable somatic or cell-free DNA contamination

SCLB permeabilizes sperm, letting DNase I fragment the genome in nearly all cells

DTT causes leakage of chromatin containing Protamine 1 from the sperm nucleus

Tn5 can access intact protamine-packed sperm chromatin without DTT treatment

## INTRODUCION

Mammalian sperm have historically been viewed primarily as vehicles for delivering paternal DNA to the oocyte at fertilization. This view is partially based on the dramatic nuclear remodeling that occurs during late spermiogenesis, in which the bulk of histones are first replaced by transition proteins and ultimately by protamines. This process generates a chromatin architecture that compacts the haploid genome into a volume approximately 10-fold smaller than that of a mature lymphocyte after taking into account the difference in ploidy. The resulting structure contains protamines cross-linked by intermolecular disulfide bonds, is transcriptionally inert and has been considered epigenetically uninformative with the exception of DNA methylation ^1,2^.

This view has been substantially revised over the past two decades ^3–5^. A small but functionally significant fraction of the sperm genome, estimated at 8-15% in mouse and approximately 5-15% in human, depending on the study, retains nucleosomal organization ^6^. Critically, these retained nucleosomes are not randomly distributed but rather they are concentrated at sites with regulatory significance for embryonic and adult development, including gene promoters and enhancers. These observations suggest that nucleosomal retention may represent an evolved mechanism for paternal transmission of regulatory information beyond the DNA sequence itself ^7–9^. A growing body of work using diverse genomic approaches has begun to characterize the organization of these retained nucleosomes. ChIP-seq studies have demonstrated that sperm nucleosomes carry histone modifications associated with both active and repressive states, including the active H3K4me3, H3K27ac, H3K4me1, the Polycomb-associated repressive modification H3K27me3, and the heterochromatin-associated H3K9me3 ^7–10^. ATAC-seq and MNase-seq analyses have mapped accessible chromatin and nucleosomal positioning across the sperm genome, identifying patterns consistent with regulatory activity after fertilization ^8,10–16^. Sperm DNA is also bound by sequence-specific transcription factors, including nuclear hormone receptors such as the estrogen and androgen receptors, as well as the architectural protein CTCF and the cohesin complex ^10,17,18^. The presence of CTCF and cohesin is particularly interesting because these factors are responsible for establishing the three-dimensional organization of the genome in somatic cells. Hi-C analyses of sperm chromatin suggest that the sperm genome is folded into chromatin loops and compartmental domains resembling those observed in somatic cells ^10,19,20^. The sperm genome is globally hypermethylated at most CpG sites compared to somatic tissues, with notable exceptions at CpG islands, which remain largely unmethylated, and at germline-specific differentially methylated regions including the paternally methylated imprinting control regions. This methylation pattern is established progressively during spermatogenesis and is largely stable from spermatogenic stages through cauda epididymal sperm ^21–23^. Together, these findings paint a picture of the sperm chromatin landscape as a structured and functionally organized epigenetic substrate rather than an inert protamine matrix, and they have direct implications for our understanding of how paternal exposures and experiences may be transmitted to offspring through non-genetic mechanisms. The biological significance of these observations has been reinforced by an expanding literature on paternal contributions to offspring phenotype. Paternal diet, stress, exercise, toxicant exposure, and metabolic state have all been shown to influence offspring physiology in mammalian models ^24–27^. Although small RNAs in sperm have received considerable attention as potential mediators of these effects, the retained nucleosomal compartment of sperm chromatin represents an additional and potentially complementary route by which paternal information could be transmitted across generations ^28^. Therefore, understanding the precise organization and architecture of sperm chromatin is of central importance to realizing the relative contribution of genes and the environment to the inheritance of phenotypes in mammals.

These advances have been made possible by the application of genomic methods, originally developed for somatic cells, to mature sperm. Mature spermatozoa in the mouse are typically isolated from the cauda epididymis by tissue dissection followed by a swim-up procedure that enriches for motile sperm ^29^. This approach yields highly purified sperm populations under standard conditions. Several investigators have additionally adopted treatments with somatic cell lysis buffer (SCLB), which contains detergents such as SDS and Triton X-100, followed by treatment with DNase I, to digest the DNA from potentially contaminating somatic cells ^30^. This approach is based on the assumption that this treatment selectively disrupts contaminating somatic cells while leaving sperm intact. The rationale for this purification step derives in part from the observation that sperm preparations from the proximal caput epididymis can be contaminated with extracellular somatic cell DNA. Sperm isolated from the caput epididymis, in particular, have been shown to be associated with web-like structures of extracellular DNA, presumably derived from the somatic compartment of the epididymal epithelium, and this contamination accounts for approximately 20–30% of cytosine methylation signal in caput sperm preparations ^31^. Critically, this contamination is largely absent or substantially reduced in sperm from the cauda epididymis, the more distal compartment from which sperm are typically harvested for chromatin analyses ^31^. Cauda sperm preparations are largely free of intact somatic cell contamination by direct microscopic examination, and standard purification methods including the swim-up procedure produce sperm samples in which somatic cell contamination represents less than one in 1000 nuclei observed by microscopy in sperm preparations ^10^. This level of contamination is unlikely to affect the results of analyses of sperm chromatin organization using genome-wide techniques such as ChIP-seq, ATAC-seq, or Hi-C.

Against this background, a recent study by Yin et al. ^32^ has proposed that cauda sperm preparations harbor approximately 2–5% contamination by somatic cell chromatin, and that this contamination is sufficient to dominate genome-wide chromatin profiles obtained by ATAC-seq, ChIP-seq, Hi-C, or similar techniques. Based on this argument, the authors conclude that previously published ATAC-seq, MNase-seq, and related analyses of mouse sperm chromatin, performed without prior treatment of sperm with somatic cell lysis buffer and DNase I, primarily reflect the chromatin organization of contaminating cell-free DNA rather than the bona fide sperm genome. The authors propose that valid sperm chromatin profiles can only be obtained following SCLB and DNase I pretreatment, supplemented by treatment with elevated concentrations of dithiothreitol (DTT) to reduce protamine disulfide bonds and permit Tn5 transposase access to sperm chromatin. If correct, this argument would imply that a substantial fraction of the published literature on mammalian sperm chromatin organization rests on flawed methodology and would require systematic re-examination. These conclusions raise important methodological concerns that warrant careful evaluation.

These concerns motivated us to revisit the question of somatic contamination in cauda sperm preparations and to characterize directly the effects of SCLB, DNase I, and DTT treatment on sperm nuclear integrity and chromatin organization. Here, we assess the extent of somatic cell and cell-free DNA contamination in cauda sperm preparations isolated by standard procedures. We then evaluate whether SCLB, used under conditions reported in the literature, selectively disrupts somatic cells without affecting sperm. We further examine the effects of DTT-mediated disulfide reduction on the integrity of the sperm nucleus and on the localization of sperm chromatin. The results indicate that properly purified cauda sperm preparations are not detectably contaminated by somatic cell DNA, and that the SCLB-DNase-DTT pretreatment is not a benign purification protocol but rather a sequence of treatments that progressively compromise sperm nuclear integrity and chromatin structure. Our findings directly contradict the methodological claims of Yin et al. ^32^ and have important implications for the interpretation of sperm chromatin profiles in published studies.

## RESULTS

### Properly purified cauda sperm preparations show no detectable contamination but subsequent treatments disrupt sperm chromatin

To examine the question of somatic cell and cell-free DNA contamination in epididymal sperm preparations, we isolated sperm from the caput and cauda epididymis of adult male mice using our previously established procedure ^10,17^. After dissection of surrounding blood vessels and fat, the epididymis was punctured with a needle, and sperm were allowed to swim up in Donner’s medium. This approach differs from that of Yin et al. ^32^ in two methodologically important aspects that may explain the observed differences in the purity of sperm preparations. First, we puncture the epididymal tubule rather than cutting the tissue into small pieces, the latter procedure being more likely to release somatic cells from the epididymal epithelium into the sperm preparation. Second, we perform the swim-up step using 15 ml tubes with 10 ml of Donner’s medium instead of 1.5 ml tubes used by Yin et al, allowing sperm to travel longer distances and better separate from potential contaminants. Using this approach, we have previously documented routine sperm preparations of at least 99.9% purity, with fewer than one somatic cell observed per 1,000 sperm counted ^10,17^.

Sperm preparations from the caput epididymis displayed the previously reported ^31^ large Hoechst-positive structures that have been attributed to extracellular DNA released from somatic cells of the epididymal epithelium. These cells can be easily distinguished from sperm by their size and shape (Figure S1A-C). Consistent with these structures representing cell-free chromatin, treatment of caput sperm preparations with DNase I eliminated them (Figure S1D-F). In contrast, sperm preparations from the cauda epididymis showed no detectable Hoechst-positive contaminating structures (Figure S1G), and these structures remained undetectable even when the image was overexposed under conditions that would have revealed weakly stained material (Figure S1H). The absence of detectable contaminating chromatin in these samples is a strong indication that, when properly purified, the contamination concern applies primarily to caput rather than cauda sperm preparations.

Although Yin et al. did not provide direct microscopic evidence for contamination of the cauda sperm preparations in their experiments, they applied incubation with SCLB followed by DNase I treatment as a standard purification step prior to chromatin analysis. The rationale for this treatment is that SCLB selectively disrupts somatic cells while leaving sperm intact, allowing DNase I to subsequently degrade any contaminating somatic DNA without affecting the sperm genome. Several previous studies have, however, reported that this treatment affects sperm beyond the elimination of putative contamination, including alterations in small RNA representation, reduced overall RNA yield, biases in detectable RNA populations, and selective loss of mitochondrial and mid-piece transcripts ^33–35^. These observations raise the possibility that SCLB-DNase I treatment compromises sperm cellular integrity more broadly than has been generally appreciated.

To examine directly whether this treatment also affects sperm chromatin, we subjected cauda sperm preparations to the individual components of the SCLB-DNase I-DTT protocol, which were performed under the same buffer compositions, concentrations, and incubation times reported by Yin et al. We first assessed nuclear integrity by Hoechst staining and microscopy (Figure 1A). Treatment with DNase I alone produced no detectable change in sperm nuclear morphology compared with untreated controls (Figure 1B), consistent with DNase I being unable to access the protamine-condensed sperm genome under standard conditions. Incubation with SCLB alone similarly produced no visible alteration in sperm nuclear structure (Figure 1C). However, the combination of SCLB incubation followed by DNase I treatment produced visibly distorted sperm head structures in approximately 3% of the sperm population (Figure 1D, Figure S1I). This 3% should be understood as a lower bound on damage rather than an estimate of its total extent, since it represents only the fraction of sperm in which structural alteration can be detected by fluorescence microscopy of Hoechst-stained material, and does not exclude additional damage to sperm chromatin in morphologically intact-appearing nuclei that would not be detected by this assay. The observed structural disruption indicates that SCLB exposure renders at least a subset of sperm permeable to DNase I, allowing the nuclease to access and degrade the sperm genome itself rather than only contaminating DNA.

**Figure 1.**
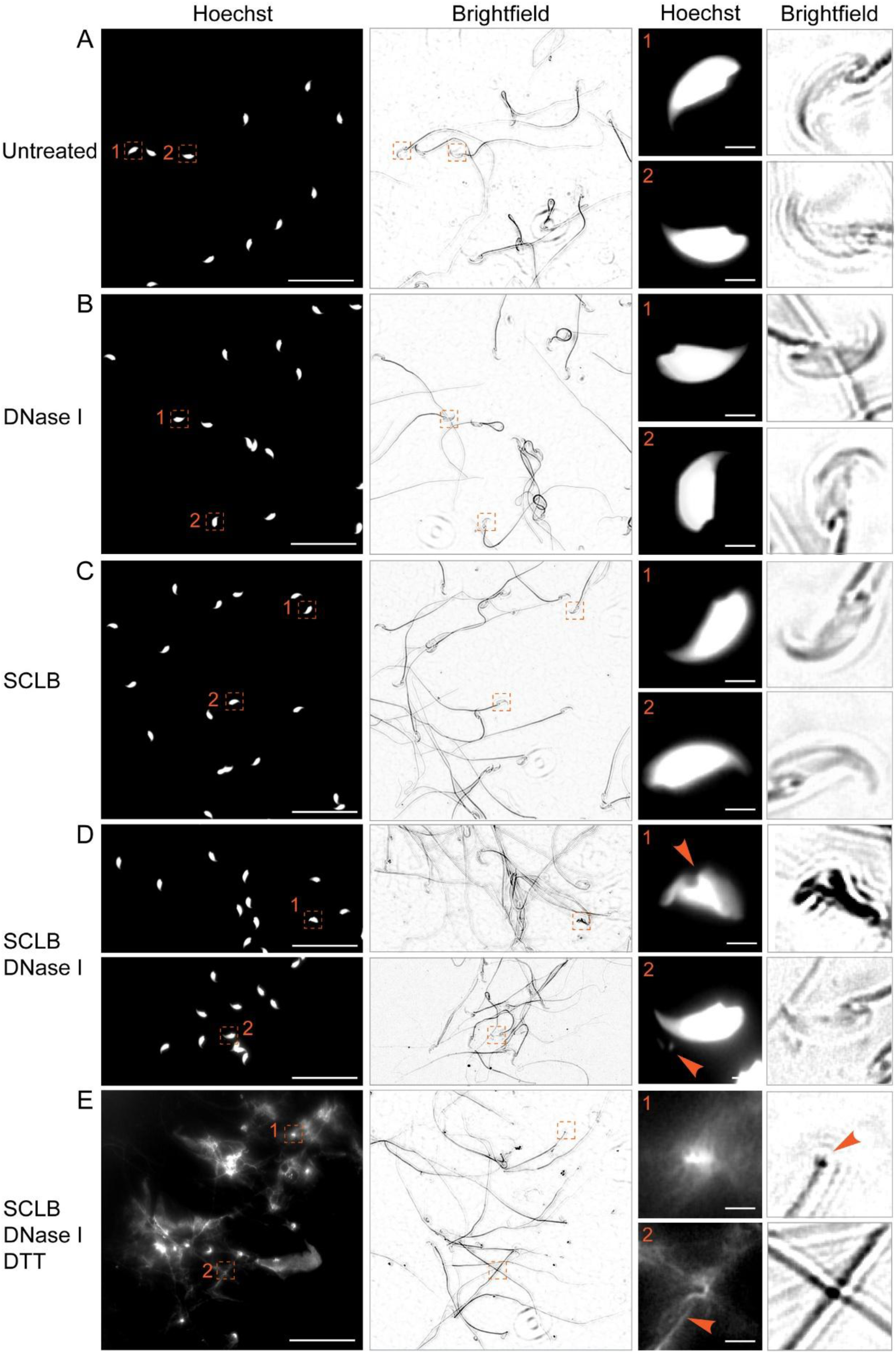
Effect of SCLB, DNase I and DTT on the integrity of the sperm nucleus. (A) Fluorescence and brightfield microscopy of mouse cauda sperm purified by the swim-up procedure. Sperm heads labeled as 1 and 2 in the left panel are shown at higher magnification on the right panels. Scale bar on left panel is 50 μm and on the right panels is 3 μm. (B) Fluorescence and brightfield microscopy of mouse cauda sperm purified as in (A) followed by treatment with DNase I. Sperm heads labeled as 1 and 2 in the left panel are shown at higher magnification on the right panels. Scale bar on left panel is 50 μm and on the right panels is 3 μm. (C) Fluorescence and brightfield microscopy of mouse cauda sperm purified as in (A) followed by treatment with SCLB. Sperm heads labeled as 1 and 2 in the left panel are shown at higher magnification on the right panels. Scale bar on left panel is 50 μm and on the right panels is 3 μm. (D) Fluorescence and brightfield microscopy of mouse cauda sperm purified as in (A) followed by treatment with SCLB and then DNase I. Sperm heads labeled as 1 and 2 in the two left panels are shown at higher magnification on the right panels. Arrowheads point to disrupted nuclear structure (1) and small pieces of Hoechst-positive nuclear material (2). Scale bar on left panel is 50 μm and on the right panels is 3 μm. (E) Fluorescence and brightfield microscopy of mouse cauda sperm purified as in (A) followed by treatment with SCLB, then DNase I, and finally DTT. Sperm heads labeled as 1 and 2 in the left panel are shown at higher magnification on the right panels. Scale bar on left panel is 50 μm and on the right panels is 3 μm. See also Figure S1

A further step in the Yin et al. protocol is treatment with DTT at high concentration to reduce intermolecular disulfide bonds between protamines, with the stated purpose of relaxing sperm chromatin sufficiently to permit access by the Tn5 transposase used in ATAC-seq. We found that DTT treatment of SCLB-DNase I-treated sperm had a substantially more severe effect on nuclear integrity than the preceding steps. Compared with the nuclear morphology of cauda sperm treated with SCLB plus DNase I (Figure 1D), part of the Hoechst-stained DNA in DTT-treated samples was no longer contained within the sperm nucleus but instead extruded from the nuclear envelope and formed an extended fibrillar network surrounding the sperm head (Figure 1E). The chromatin was thus released from its native nuclear context, and this affects 100% of all sperm cells present in the preparation.

Together, these results indicate that cauda sperm preparations obtained by standard procedures are not detectably contaminated by somatic-cell or cell-free DNA in sufficiently high amounts that can be observed by DNA staining followed by fluorescence microscopy, that the SCLB-DNase I treatment proposed to address putative contamination instead compromises a fraction of sperm nuclei and exposes the sperm genome to nuclease degradation. Furthermore, subsequent DTT treatment disrupts the residual nuclear architecture and releases part of the sperm chromatin from the nucleus. We further note that the protocol as published requires multiple washes and centrifugation steps between treatments, with centrifugation conditions that were not reported by Yin et al. and may further contribute to sperm chromatin alterations.

### Treatment with SCLB-DNase I and DTT progressively degrades the sperm genome and produces DNA damage within the sperm nucleus

The disruption of sperm nuclear architecture visualized by fluorescence microscopy suggests that treatments involving SCLB-DNase I and DTT allow nuclease access to the sperm genome and disruption of nuclear architecture. To test this directly, we isolated genomic DNA from cauda sperm subjected to each step of the process and assessed DNA integrity by gel electrophoresis (Figure 2A). DNA from untreated sperm migrated predominantly as a single high-molecular-weight band of approximately 30 kb, with minimal signal at lower molecular weights. Treatment with DNase I alone produced only an apparent slight reduction in the size of the main band, consistent with DNase I being largely unable to access the protamine-condensed sperm genome in the absence of permeabilization (Figure 2A). Incubation with SCLB followed by DNase I treatment, however, resulted in substantial degradation of the sperm genome, with a marked smear of lower-molecular-weight DNA below the main band. Further treatment with DTT produced still more extensive degradation, with progressively more of the genomic DNA migrating as low-molecular-weight fragments (Figure 2A).

**Figure 2.**
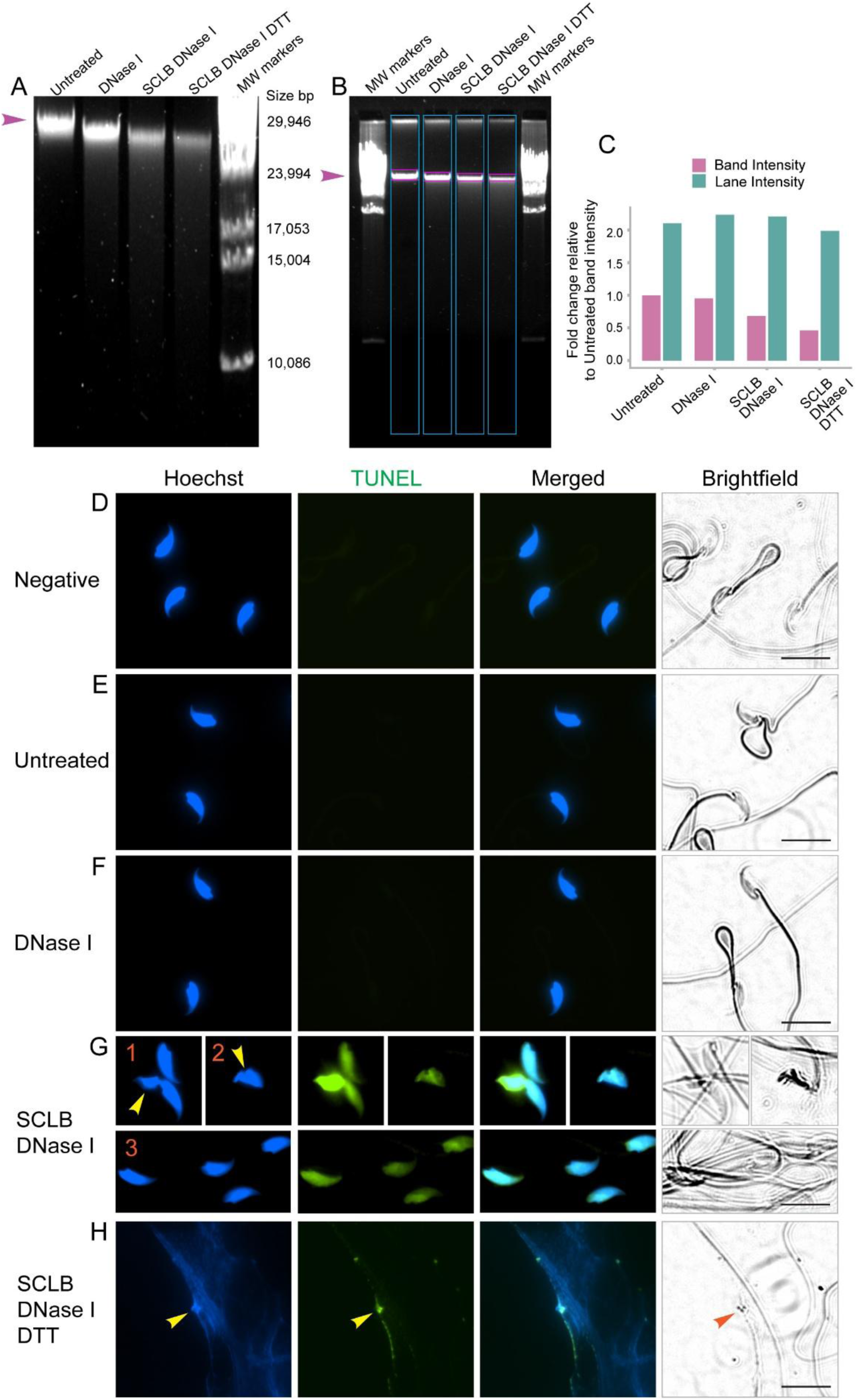
Effects of SCLB, DNase I and DTT on the integrity of sperm DNA. (A) Agarose gel electrophoresis of DNA isolated from cauda sperm purified by the swim-up procedure and subjected to the treatments indicated above. (B) Agarose gel electrophoresis of DNA isolated from sperm purified by the swim-up procedure and subjected to the treatments indicated above. Boxes in teal and purple indicate areas where DAPI staining was quantified in the results shown in panel (C). (C) Quantification of areas in the agarose gel shown in panel (B). Colors correspond to those used in the boxes shown in (B). (D) Results from TUNEL assay and fluorescence microscopy of purified mouse sperm. Negative result without TUNEL assay components. Scale bars correspond to 13 μm. (E) Results from TUNEL assay and fluorescence microscopy of purified mouse sperm. Untreated sperm. Scale bars correspond to 13 μm. (F) Results from TUNEL assay and fluorescence microscopy of purified mouse sperm. Sperm treated with DNase I only. Scale bars correspond to 13 μm. (G) Results from TUNEL assay and fluorescence microscopy of purified mouse sperm. Sperm treated with SCLB followed by DNase I. Arrowheads point to defective sperm heads based on Hoechst staining. Scale bars correspond to 13 μm. (H) Results from TUNEL assay and fluorescence microscopy of purified mouse sperm. Sperm treated with SCLB followed by DNase I and then DTT. Arrowhead points to defective sperm head surrounded by leaked DNA. Scale bars correspond to 13 μm. See also Figure S2

Quantification of the gel signal allowed us to assess the extent of degradation in two complementary ways. The absolute signal in the high-molecular-weight band decreased progressively with each step of the protocol relative to untreated sperm, indicating loss of intact DNA (Figures 2B and 2C). Because absolute band intensity can be influenced by loading variation and DNA recovery efficiency, we also measured the fraction of DNA in the main band relative to the total DNA signal in each lane (Figure 2B). The total signal per lane was comparable across treatments, indicating that the DNA was retained in the samples rather than lost during isolation, but the fraction present as intact high-molecular-weight DNA decreased progressively with each step (Figure 2C). The SCLB-DNase I treatment with or without DTT therefore results in extensive fragmentation of the sperm genome itself.

To confirm that the observed DNA fragmentation reflects nuclease activity within the sperm nucleus rather than damage occurring during DNA isolation, we used TUNEL assays to localize 3′-hydroxyl ends, which can be produced by DNase I digestion, within sperm cells. Untreated cauda sperm and sperm treated with DNase I alone showed no detectable TUNEL signal within the limits of the assay (Figures 2D-F; Figures S2A-C), consistent with the gel data and with sperm chromatin being inaccessible to DNase I in untreated sperm. Sperm incubated with SCLB and then treated with DNase I, in contrast, displayed clear TUNEL signal within the nucleus in virtually all sperm cells (Figure 2G; Figure S2D), suggesting that SCLB renders the nuclear interior accessible to DNase I and that the resulting DNA cleavage occurs in situ rather than during downstream processing. Further treatment with DTT produced TUNEL signal not only within but also surrounding the sperm head, in the leaked fibrillar chromatin documented in Figure 1E, with damage extending throughout this released material (Figure 2H and Figure S2E). Quantification of TUNEL signal across the treatment conditions confirms and extends the qualitative observations (Figure S2F, S2G). The negative control, untreated cauda sperm, and sperm treated with DNase I alone all showed statistically indistinguishable TUNEL signal at baseline, confirming that DNase I cannot access intact sperm chromatin. In contrast, mean TUNEL signal intensity per sperm increased approximately 2.5-fold in samples treated with SCLB followed by DNase I. This quantification could not be performed in samples treated with DTT due to the difficulty in assigning specific DNA webs to individual sperm heads.

Together, these observations indicate that the morphologically disrupted population identified microscopically with Hoechst staining after SCLB-DNase I treatment represents only those sperm in which damage has progressed sufficiently to alter the visible architecture of the sperm head, while virtually all sperm has sustained DNA damage without showing visible morphological change. These findings establish that the SCLB-DNase I-DTT protocol applied to cauda sperm preparations does not selectively eliminate contaminating somatic DNA but instead progressively damages the sperm genome itself. Chromatin analyses performed on sperm subjected to this protocol therefore reflect material that has been fragmented, denucleated, and structurally rearranged by the treatments involved.

### Treatment with SCLB, DNase I and DTT disrupts the localization of sperm chromatin proteins

The findings presented above indicate that the SCLB-DNase I-DTT treatment fragments the sperm genome and extrudes chromatin from the nucleus. Beyond these effects on the DNA itself, the protocol may also disrupt the interactions of nuclear proteins with sperm chromatin. Treatment with 50 mM DTT, in particular, is designed to reduce the disulfide bonds that cross-link protamines, but at this concentration DTT can also affect other chromatin proteins through several mechanisms, including reduction of redox-sensitive cysteines in DNA-binding domains, weak chelation of Zn²⁺ through the two thiol groups of DTT, and consequent destabilization of zinc finger protein-DNA interactions ^36–39^. We therefore examined the consequences of the SCLB-DNase I-DTT treatments on the nuclear localization of three structurally important sperm chromatin proteins: Protamine 1, the major protein of sperm chromatin condensation; CTCF, the principal architectural factor establishing chromatin loops in somatic cells; and SMC1, a core subunit of the cohesin complex that cooperates with CTCF in three-dimensional genome organization.

We first examined the distribution of Protamine 1 using monoclonal antibodies with established high specificity for this protein ^40^. In untreated cauda sperm, Protamine 1 was detected by immunofluorescence microscopy in two locations. Protamine 1 is located throughout the sperm nucleus, as expected for the major chromatin compaction protein and, surprisingly, in the midpiece of the sperm tail (Figure 3A; Figure S3A). To confirm that the midpiece signal reflects genuine intracellular localization of Protamine 1 rather than non-specific surface association, we performed confocal z-stack imaging and reconstructed both longitudinal (Figure S3F) and transverse (cross-sectional) (Figure S3G) views of the sperm mid-piece tail. In longitudinal z-stack sections, the Protamine 1 signal persisted continuously across consecutive optical slices throughout the entire depth of the midpiece (z slices = 4 - 16). In transverse sections, if the signal originated from proteins bound to the exterior of the sperm membrane, fluorescence signal would be expected to appear in the top peripheral slice (e.g. z = 4), disappear in the central slice (e.g. z = 10) and reappear in the bottom peripheral slice (e.g. z= 16). Figure S3G represents a cross-sectional reconstruction generated from the same serial z-stack sections shown in Figure S3F at the same yellow reference cross. This reconstitution compiles the consecutive optical slices across the z-axis to visualize the fluorescence distribution through the diameter of the sperm tail. Figure S3G confirms that the signal is not localized at the periphery of the tail but remains centered within the midpiece, supporting an internal luminal localization of Protamine 1. It is possible that this cytoplasmic Protamine 1 protein is a remnant left during the spermiogenesis process.

**Figure 3.**
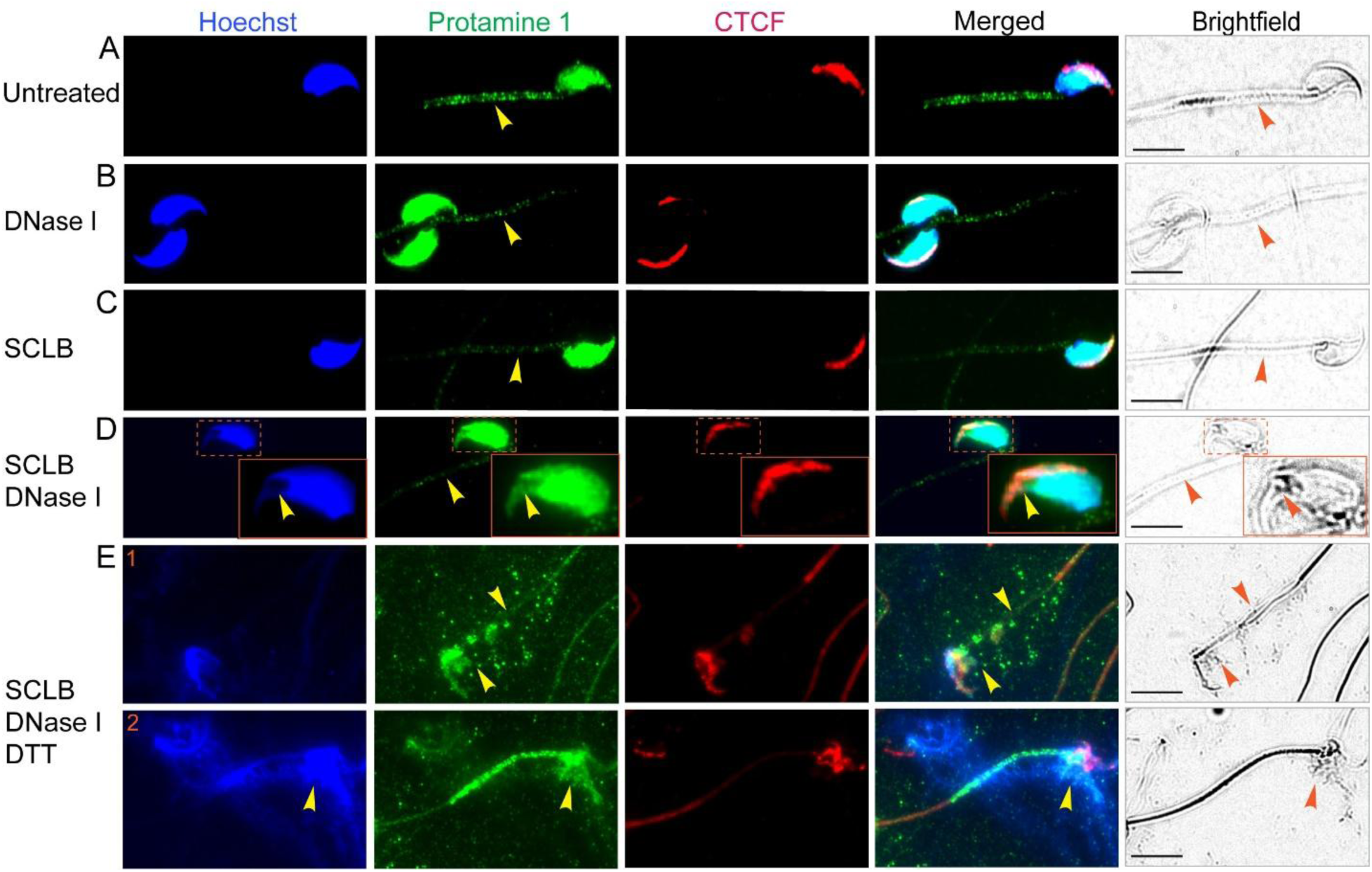
Effects of SCLB, DNase I and DTT on the nuclear distribution of Protamine 1 and CTCF. (A) Immunofluorescence microscopy of purified untreated cauda sperm using antibodies to Protamine 1 and to CTCF. Arrowhead points to Protamine 1 staining in the midpiece of the sperm tail. Scale bars correspond to 7 μm (B) Immunofluorescence microscopy of purified cauda sperm treated with DNase I using antibodies to Protamine 1 and to CTCF. Arrowhead points to Protamine 1 staining in the midpiece of the sperm tail. Scale bars correspond to 7 μm. (C) Immunofluorescence microscopy of purified cauda sperm treated with SCLB using antibodies to Protamine 1 and to CTCF. Arrowhead points to Protamine 1 staining in the midpiece of the sperm tail. Scale bars correspond to 7 μm. (D) Immunofluorescence microscopy of purified cauda sperm treated with SCLB followed by DNase I using antibodies to Protamine 1 and to CTCF. Arrowhead points to Protamine 1 staining in the midpiece of the sperm tail. A second arrowhead points to defects in the sperm nucleus. Scale bars correspond to 7 μm. (E) Immunofluorescence microscopy of purified cauda sperm treated with SCLB followed by DNase I, and then DTT using antibodies to Protamine 1 and to CTCF. Arrowhead points to a misshapen sperm head caused by leakage of sperm DNA. Scale bars correspond to 7 μm. See also Figure S3

DNase I treatment alone did not visibly alter Protamine 1 distribution (Figure 3B; Figure S3B). However, treatment with SCLB, either alone or followed by DNase I, decreased the midpiece signal while preserving nuclear localization at this level of resolution (Figure 3C-D; Figure S3C-D). This pattern suggests that SCLB-induced permeabilization of the sperm plasma membrane allows the Protamine 1 pool in the midpiece to exit the cell, while the heavily cross-linked nuclear pool remains in place, although the treatment alters the nuclear integrity of some sperm nuclei (Figure 2D). Further treatment with DTT produced a markedly different pattern: Protamine 1 is no longer confined to the sperm nucleus but instead remains associated with the DNA that has leaked into the surrounding milieu (Figure 3E; Figure S3E), consistent with disulfide reduction releasing protamine-protamine interactions while preserving the underlying protamine-DNA association. Unexpectedly, Protamine 1 also reappeared in the sperm midpiece after DTT treatment (Figure 3E; Figure S3E), perhaps reflecting association of the protein itself or in combination with DNA to the surface of the sperm (Figure 3E2, Hoechst panel).

We next examined the distribution of CTCF, which is critical to the question of whether the SCLB-DNase I-DTT protocol preserves the three-dimensional chromatin organization that this protein establishes. Prior work has demonstrated that sperm chromatin is organized into loops and compartmental domains broadly resembling those of somatic cells, with CTCF and cohesin playing analogous roles to those they perform in other cell types ^10,17,19^. In contrast, Yin et al. reported that sperm chromatin treated with the SCLB-DNase I-DTT protocol lacks the standard contact domains observed in somatic cells and is instead highly disorganized. The DNA eviction from the nucleus after DTT treatment documented in Figures 1 and 2 above provides one likely explanation for this finding, since chromatin released from the nucleus into the surrounding milieu would not retain the spatial organization characteristic of an intact nucleus, regardless of whether the underlying protein-DNA interactions remained intact. An additional possibility is that the treatments themselves disrupt the binding of architectural proteins to DNA, which could compromise three-dimensional organization independently of nuclear removal. Immunofluorescence microscopy with antibodies against CTCF shows that treatment with SCLB, DNase I, or the combination of the two did not detectably alter CTCF nuclear localization at the resolution afforded by this approach (Figures 3A-D and Figures S3A-D). After additional DTT treatment, CTCF appears to stay bound to DNA, and this DNA remains associated with the nucleus or the sperm head rather than the DNA web-like structures released from the nucleus (Figures 3E and Figure S3E).

We also examined the distribution of SMC1, a core subunit of the cohesin complex that cooperates with CTCF in establishing three-dimensional chromatin architecture. In untreated cauda sperm, SMC1 was distributed throughout the nucleus in a pattern that overlaps with CTCF but extends over a broader spatial domain, consistent with the presence of cohesin at genomic sites that lack CTCF observed in somatic cells ^41^. As with Protamine 1, SMC1 was also detected in the sperm midpiece at what appear to be very high levels (Figure 4A; Figure S4A). Confocal cross-sectional analysis confirmed that the midpiece SMC1 signal reflected genuine intracellular localization rather than non-specific binding to the sperm membrane (Figure S4F, S4G). DNase I treatment alone did not alter SMC1 distribution (Figure 4B; Figure S4B). Treatment with SCLB caused loss of SMC1 from the midpiece, consistent with the membrane permeabilization that also released Protamine 1 from this compartment (Figure 4C; Figure S4C). The combination of SCLB followed by DNase I does not appear to affect the nuclear localization of SMC1 (Figure 4D; Figure S4D). Subsequent DTT treatment results in SMC1 remaining in or in close association with the nucleus or the sperm head (Figure 4E; Figure S4E), as was also observed for CTCF.

**Figure 4.**
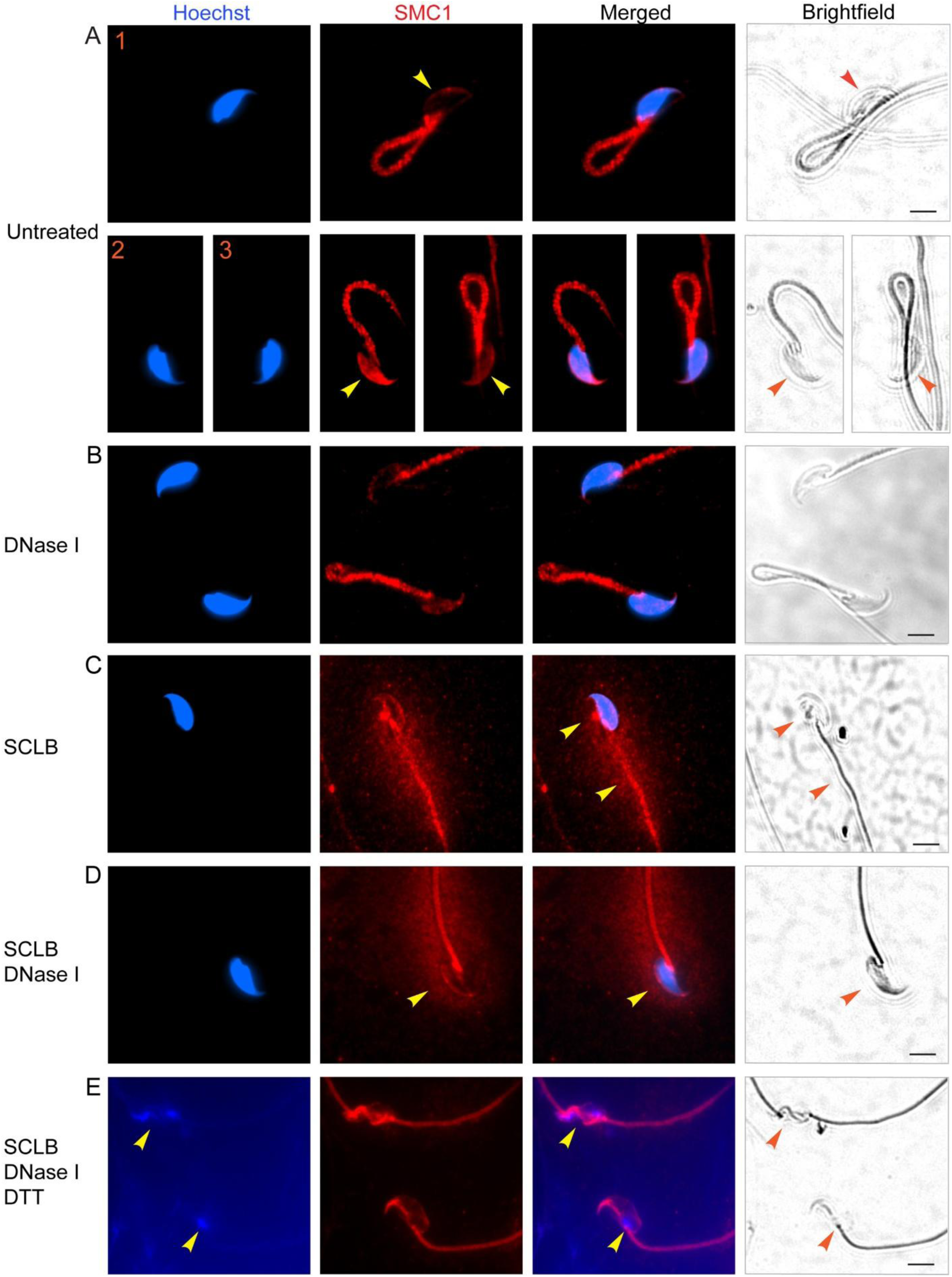
Effects of SCLB, DNase I and DTT on the nuclear distribution of cohesin. (A) Immunofluorescence microscopy of purified untreated cauda sperm using antibodies to SMC1. Arrowhead points to nuclear staining of SMC1, which ranges from low (panel 1) to high (panel 2) or intermediate (panel 3). SMC1 shows very strong staining in the midpiece of the sperm tail. Scale bars correspond to 30 μm. (B) Immunofluorescence microscopy of purified cauda sperm treated with DNase I using antibodies to SMC1. Scale bars correspond to 30 μm. (C) Immunofluorescence microscopy of purified cauda sperm treated with SCLB using antibodies to SMC1. Arrowhead points to SMC1 staining that appears to emanate from the midpiece of the sperm tail. Scale bars correspond to 30 μm. (D) Immunofluorescence microscopy of purified cauda sperm treated with SCLB followed by DNase I using antibodies to SMC1. Arrowhead points to SMC1 staining in the outside region surrounding the midpiece of the sperm tail. Nuclear staining appears to be largely undisturbed. Scale bars correspond to 30 μm.(E) Immunofluorescence microscopy of purified cauda sperm treated with SCLB followed by DNase I, and then DTT using antibodies to SMC1. Arrowhead points to a misshapen sperm head caused by leakage of sperm DNA. SMC1 appears to be contained within the nucleus. Scale bars correspond to 30 μm. See also Figure S4

Together, these findings suggest that the SCLB-DNase I-DTT protocol affects sperm chromatin proteins differently and at different steps of the protocol. Membrane permeabilization by SCLB releases proteins from the sperm midpiece, where they may not play a functional role, without affecting nuclear localization. DTT treatment discharges some chromatin from the nucleus into the surrounding milieu, with Protamine 1 remaining variably associated with this released material whereas CTCF and SMC1 appear to remain mostly in the nuclear space. The combined effect is a progressive displacement and reorganization of the protein-DNA complexes that establish nuclear organization.

### Treatment of sperm with SCLB-DNase I-DTT affects the localization of H3K4me3

To further explore the consequences of SCLB-DNase I-DTT treatment on sperm chromatin, we examined the distribution of H3K4me3 after each treatment step using fluorescence microscopy. In untreated sperm, H3K4m3 is located in the nucleus but most of the signal is associated with the midpiece of the tail. Immunofluorescence signal can also be observed in the principal piece of the tail (Figures 5A and S5A). Incubation with DNase I, SCLB, or SCLB followed by DNase I had no apparent effect on the distribution of H3K4me3 in the sperm nucleus or tail, although the SCLB-DNase I treatment appear to affect the integrity of a subset of sperm heads (Figures 5A-D and S5A-D). Subsequent treatment with DTT, however, resulted in leakage of some of the DNA from the nucleus, as in previous experiments. Most or all H3K4me3 remained associated with the DNA retained in the nucleus (Figures 5E2 and S5E); although a fraction of H3K4me3 appears to be present in the leaked material, it is in fact associated with the principal piece of the sperm tail rather than leaked chromatin (Figures 5E and S5E). These results confirm our observations above suggesting that DTT treatment allows chromatin to escape the nucleus, severely disrupting nuclear organization.

**Figure 5.**
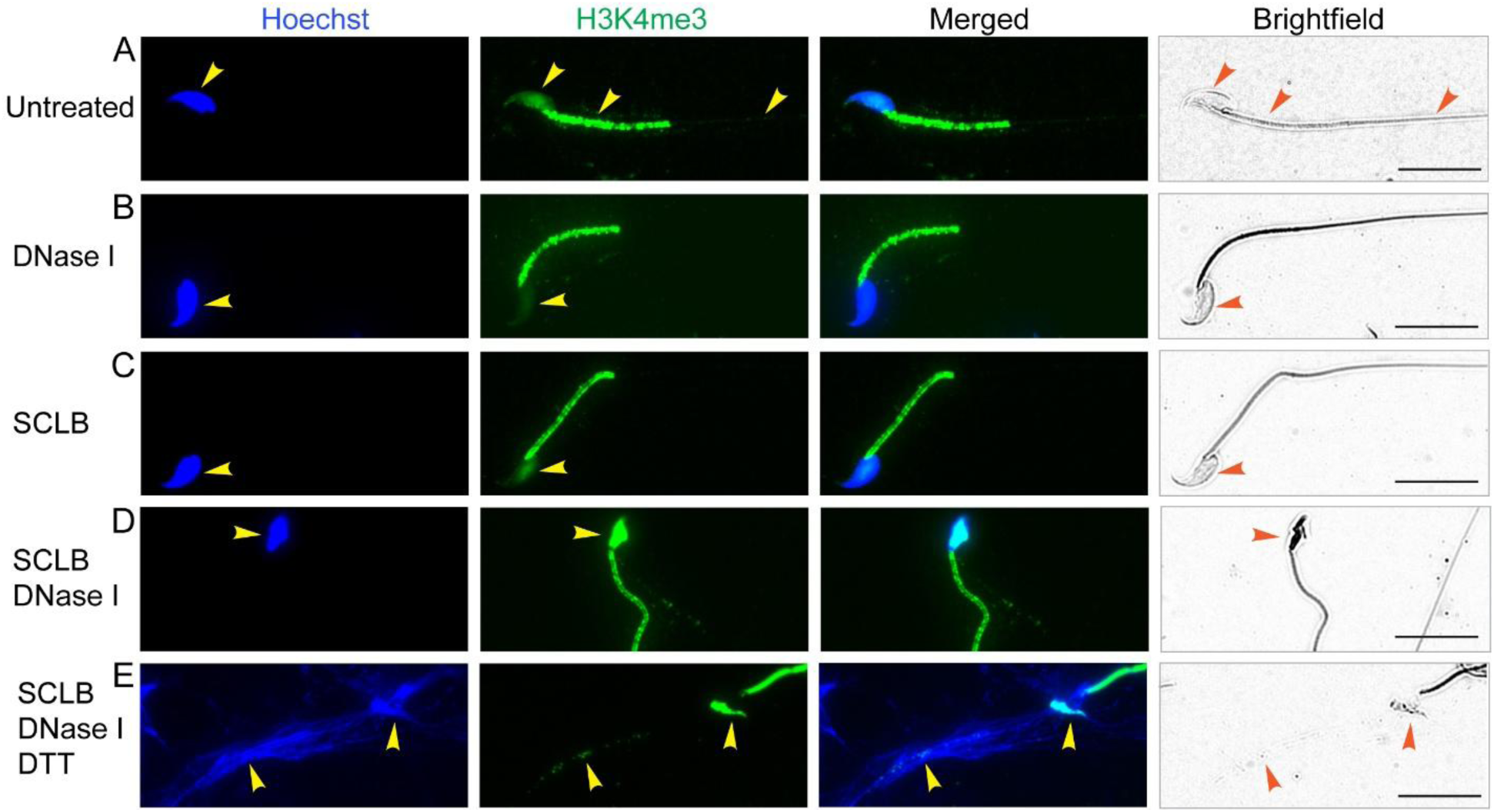
Effects of SCLB, DNase I and DTT on the nuclear distribution of H3K4me3. (A) Immunofluorescence microscopy of purified untreated cauda sperm using antibodies to H3K4me3. Arrowheads points to nuclear staining of H3K4me3, which also shows very strong staining in the midpiece of the sperm tail. Staining can also be observed in the principal piece of the tail. Scale bars correspond to 8 μm. (B) Immunofluorescence microscopy of purified cauda sperm treated with DNase I using antibodies to H3K4me3. Arrowhead points to nuclear staining of Hoechst or H3K4me3. Scale bars correspond to 8 μm. (C) Immunofluorescence microscopy of purified cauda sperm treated with SCLB using antibodies to H3K4me3. Arrowhead points to nuclear staining of Hoechst or H3K4me3. Scale bars correspond to 8 μm). (D) Immunofluorescence microscopy of purified cauda sperm treated with SCLB followed by DNase I using antibodies to H3K4me3. Arrowhead points to nuclear staining of Hoechst or H3K4me3 in a distorted sperm head. Scale bars correspond to 8 μm. (E) Immunofluorescence microscopy of purified cauda sperm treated with SCLB followed by DNase I, and then DTT using antibodies to H3K4me3. Arrowhead points to a misshapen sperm head that contains most of the H3K4me3. A second arrowhead point to H3K4me3 signal associated with the principal piece of the sperm tail. Scale bars correspond to 8 μm. See also Figure S5

As we have observed for other chromatin proteins, H3K4me3 also associates with the midpiece of the sperm tail. To determine whether this signal reflects intracellular localization rather than non-specific surface association, we performed confocal z-stack imaging and reconstructed both longitudinal and cross-sectional views of the midpiece (Figures S5F and S5G). H3K4me3 was continuously present across consecutive optical slices throughout the entire depth of the midpiece (Figure S5F). In transverse sections, the H3K4me3 signal was centered within the midpiece rather than at the periphery, suggesting that H3K4me3 localizes to the lumen of the midpiece (Figure S5G).

### Tn5 transposase can access sperm chromatin without DTT treatment

A central rationale for the SCLB-DNase I-DTT protocol is that intact sperm chromatin is presumed to be inaccessible to the enzymatic reagents used in chromatin profiling, including restriction enzymes and the Tn5 transposase used in ATAC-seq. Under this premise, DTT-mediated reduction of protamine disulfide bonds is considered necessary to relax sperm chromatin sufficiently to permit Tn5 insertion. A direct consequence of this premise is the argument advanced by Yin et al. that ATAC-seq signal obtained from sperm preparations that have not been pretreated with DTT cannot reflect insertion into bona fide sperm chromatin and must instead reflect insertion into the chromatin of contaminating somatic cells. The validity of this argument depends critically on the assumption that Tn5 cannot access protamine-condensed sperm chromatin in the absence of DTT pretreatment. We tested this assumption directly using ATAC-see, a variation of the ATAC-seq technique in which Tn5 is loaded with fluorescently labeled transposon adapters so that successful chromatin insertion can be visualized by fluorescence microscopy.

Untreated cauda sperm incubated without Tn5 transposase showed no detectable fluorescence signal, establishing the assay baseline (Figure 6A; Figure S6A). When untreated cauda sperm were subjected to the complete ATAC-see reaction including Tn5, however, a clear fluorescent signal was observed throughout the nuclei of all sperm in the sample (Figure 6B; Figure S6B). The signal was distributed across the entire nuclear volume rather than confined to specific subnuclear regions, indicating that Tn5 had successfully accessed sperm chromatin and inserted fluorescent adapters at multiple sites genome-wide. The morphology of the labeled cells was unambiguously that of mature sperm, i.e. elongated, flattened nuclei with the characteristic falciform shape of mouse sperm heads. This morphology is readily distinguishable from any potential contaminating somatic cell, which would present a round nuclear morphology of substantially larger diameter. In addition, sperm nuclei are adjacent to sperm heads visible in brightfield microscopy images. The signal therefore originates from sperm chromatin itself, not from contaminating somatic cell chromatin. No other signal is observed in the sperm preparation outside of sperm heads, suggesting the absence of extracellular cel-free DNA of somatic cell origin. This finding contradicts the core premise on which the requirement for DTT pretreatment rests. Sperm chromatin in the cauda epididymis, despite its protamine-mediated condensation, is sufficiently accessible to Tn5 transposase to support efficient adapter insertion across the entire nuclear volume. ATAC-seq profiles obtained from untreated cauda sperm therefore cannot be interpreted as necessarily reflecting somatic cell contamination; they reflect insertion into a chromatin substrate that Tn5 can access readily.

**Figure 6.**
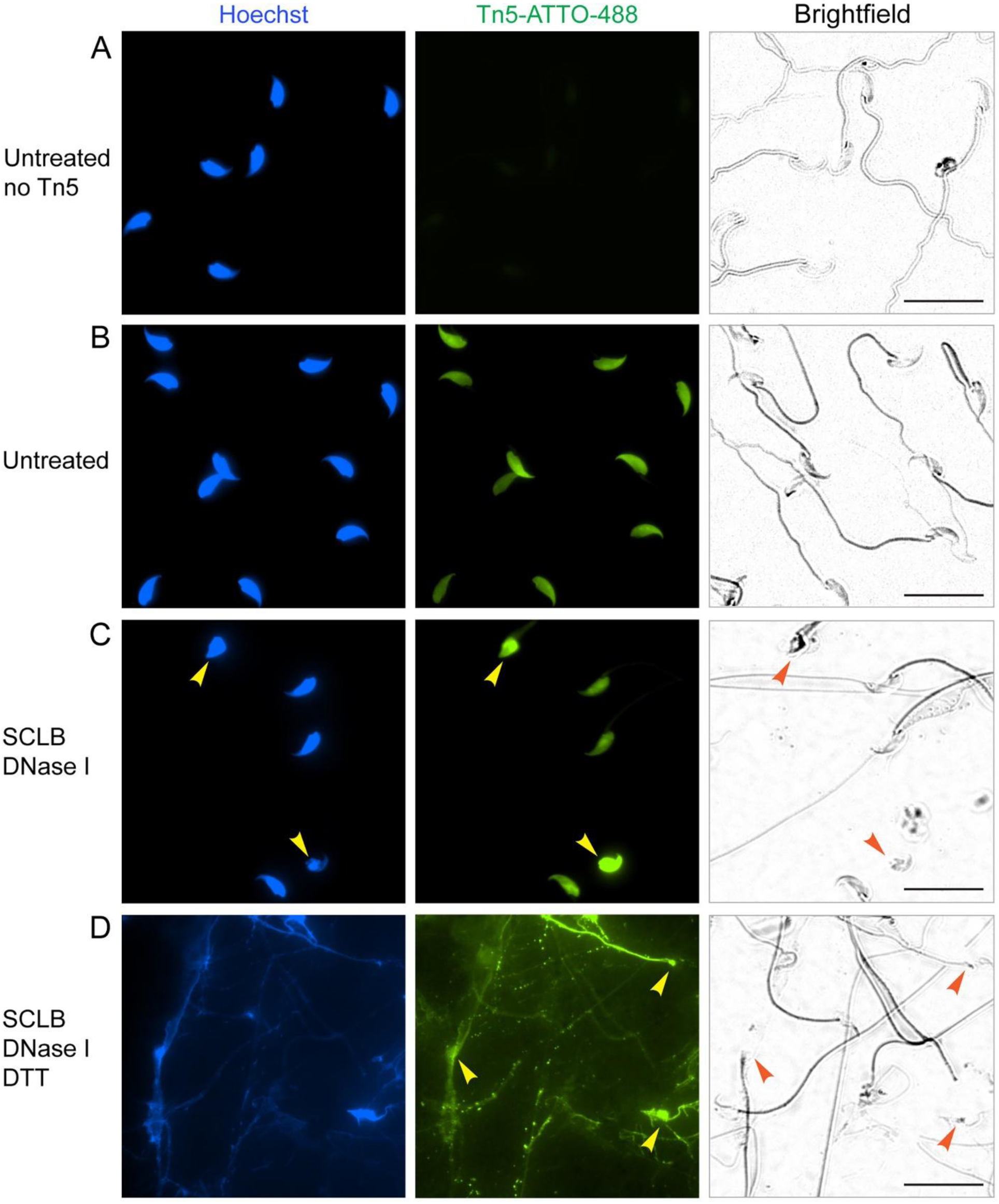
ATAC-see analysis of sperm chromatin. (A) Fluorescence microscopy of purified untreated cauda sperm subjected to the ATAC-see procedure with no Tn5 added. Scale bars correspond to 25 μm. (B) Fluorescence microscopy of purified untreated cauda sperm subjected to the ATAC-see procedure. Scale bars correspond to 25 μm. (C) Fluorescence microscopy of purified cauda sperm treated with SCLB followed by DNase I and subjected to the ATAC-see procedure. Arrowheads point to two sperm nuclei with abnormal morphology. These two nuclei have stronger ATAC-see signal. Scale bars correspond to 25 μm. (D) Fluorescence microscopy of purified cauda sperm treated with SCLB followed by DNase I and then DTT. The sperm sample was then subjected to the ATAC-see procedure. Arrowheads point to sperm heads. Scale bars correspond to 25 μm. See also Figure S6

We next examined how the components of the SCLB-DNase I-DTT protocol affect this baseline ATAC-see signal in untreated sperm. Treatment with SCLB followed by DNase I did not appreciably alter the ATAC-see signal compared with untreated sperm in most nuclei, with the labeled signal remaining distributed throughout the sperm nucleus (Figure 6C; Figure S6C). However, a subset of nuclei, identified in Figure 1D as structurally abnormal due to the treatment, appears to have a stronger signal suggestive of increase accessibility to sperm chromatin. Subsequent treatment with DTT produced a strikingly different distribution of ATAC-see signal, with the fluorescent label now found in remnants of sperm heads as well as DNA that had been discharged from the sperm nucleus (Figure 6D; Figure S6D). The DTT-treated sample therefore presents ATAC-see reactivity not only in intact sperm nuclear chromatin but also in released DNA whose spatial organization no longer corresponds to its native nuclear context. Together, these findings establish two points with direct relevance to the interpretation of published ATAC-seq data on sperm chromatin. First, Tn5 can access intact protamine-condensed sperm chromatin without prior disulfide reduction, undermining the assumption that DTT pretreatment is necessary for valid sperm ATAC-seq. ATAC-seq profiles obtained from properly purified cauda sperm without SCLB-DNase I-DTT pretreatment therefore reflect insertion into sperm chromatin and not into contaminating somatic cell chromatin. Second, the DTT step does not merely make sperm chromatin accessible to Tn5. Rather, it relocates part of the chromatin from its native nuclear context into a form whose architectural organization has been disrupted, as shown in this and preceding sections. The differences between such profiles and those obtained from untreated sperm cannot be interpreted as the difference between contaminated and decontaminated sperm chromatin.

### Accessible regions identified by ATAC-seq in untreated sperm are bona fide sites in sperm chromatin

To examine the accessibility of sperm chromatin to the Tn5 transposase during the ATAC-seq procedure, and to assess the effect of the SCLB–DNase I–DTT treatments, we performed ATAC-seq on sperm purified from the cauda epididymis. Sequencing reads were partitioned by insert size into a sub-nucleosomal fraction (50–115 bp) and a nucleosomal fraction (180–247 bp), corresponding respectively to regions protected from Tn5 insertion by transcription factors and by nucleosomes ^42^. We retained the sub-nucleosomal reads, called peaks, and defined their summits. Summits from untreated, SCLB–DNase I, and SCLB–DNase I–DTT samples were pooled and used as anchors for all subsequent analyses. In parallel, we reprocessed the ATAC-seq data of Yin et al ^32^ (GSE226685), including several samples that had been deposited but not presented in their study.

To compare our data directly with those of Yin et al, we clustered the signal at these anchors into three groups; Figures 7A and S7A show the resulting heatmaps for our samples and Figures 7B and S7B the parallel analysis of the Yin et al data. Clusters C1–C2 correspond to the accessible sites that Yin et al attributed to contaminating somatic-cell DNA, whereas cluster C3 contains the sites they assign to sperm. All three clusters are present in our untreated sperm, but the C3 sites are absent from the untreated sperm of Yin et al. Accessibility at these sites appears to be highly sensitive to the amount of Tn5. Reducing the enzyme from 2.5 µl to 1.5 µl markedly lowered accessibility across cluster C3, which may account for the failure of Yin et al to detect these sites (Figures 7C and S7C).

**Figure 7.**
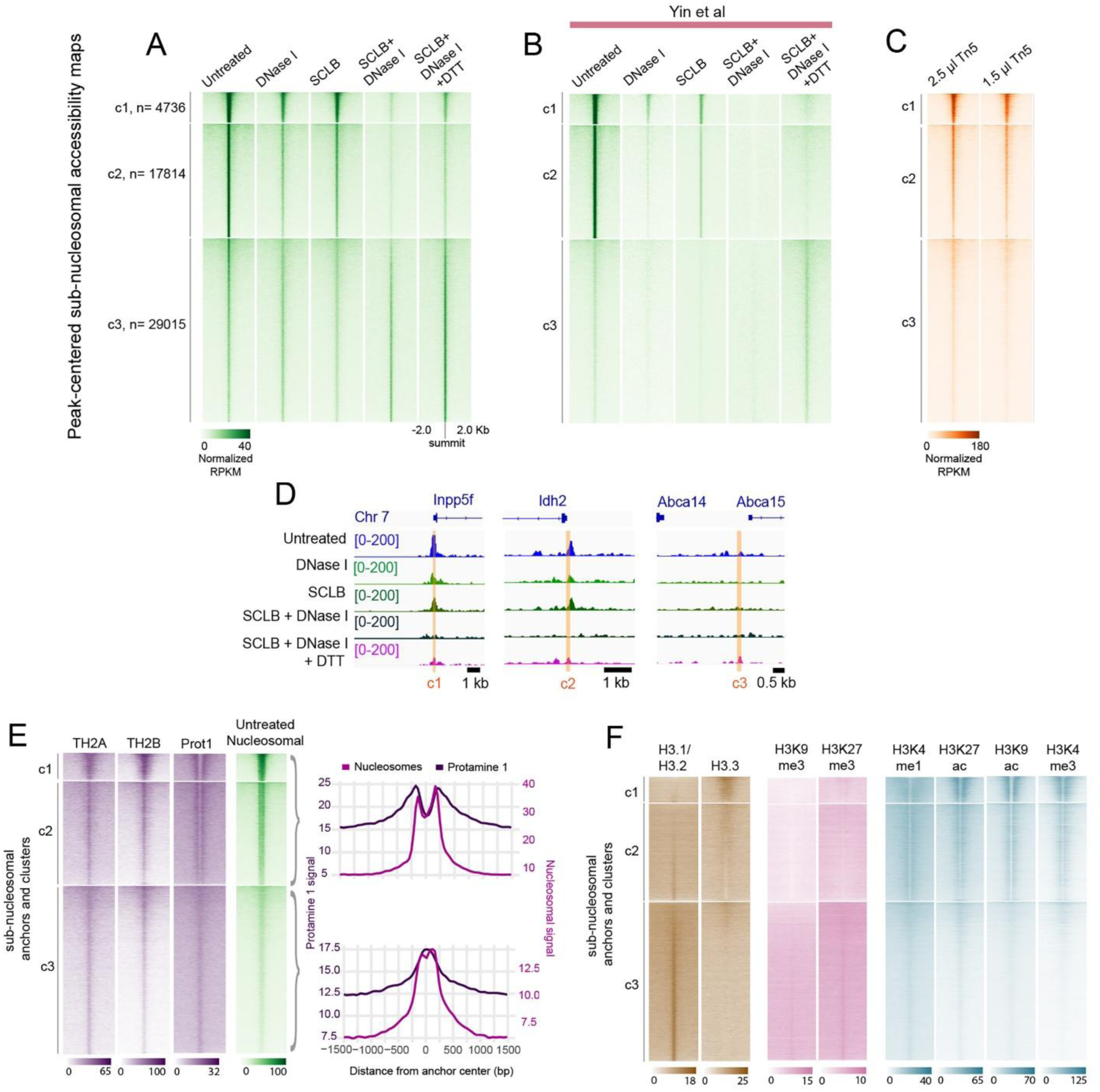
ATAC-seq analysis of sperm chromatin. (A) Heatmaps of results from ATAC-seq experiments performed with purified cauda sperm. Sperm preparations were subjected to the treatments indicated above each of the heatmaps followed by the ATAC-seq procedure. Anchors are summits of peaks of sub-nucleosome size reads. Signal corresponds to sub-nucleosome-size reads. (B) Heatmaps of results from ATAC-seq experiments performed with cauda sperm by Yin et al and downloaded from GEO accession GSE226685. Sperm preparations were subjected to the treatments indicated above each of the heatmaps followed by the ATAC-seq procedure as described by Yin et al. Data analysis was performed exactly as for the results described in panel (A). Anchors are summits of peaks of sub-nucleosome size reads. Signal corresponds to sub-nucleosome-size reads. (C) Results of ATAC-seq experiments performed with purified untreated cauda sperm using 2.5 μl and 1.5 μl of Tn5 transposase. Anchors are summits of peaks of sub-nucleosome size reads. Signal corresponds to sub-nucleosome-size reads. (D) Representative examples of ATAC-seq accessible sites from each of the c1-c3 clusters shown in panel (A). Each row indicates the treatment performed on purified cauda sperm before the ATAC-seq procedure. (E) Analysis of the distribution of ChIP-seq data for Protamine 1 and sperm-specific histones TH2A and TH2B on the c1-c3 clusters obtained in panel (A). Also shown is the location of nucleosomes in the same clusters based on the distribution of nucleosome-size reads from ATAC-seq experiments. Anchors are summits of peaks of sub-nucleosome size reads. The right-side panels show the average enrichment profiles for the nucleosome and Protamine 1 signals for clusters c1-c2 (top) and c3 (bottom). (F) Distribution of ChIP-seq data for histone H3 variants and histone covalent modifications on anchors for clusters c1-c3 defined in panel (A). Anchors are summits of peaks of sub-nucleosome size reads.

DNase I or SCLB applied individually produced only a slight change in signal intensity, most noticeably at clusters c1–c2. Because neither treatment altered DNA integrity by TUNEL, we attribute this modest effect to mechanical stress from the additional centrifugation steps these treatments require ^43^, an effect that is more pronounced in the Yin et al samples. If sites in clusters c1-c3 correspond to cell-free DNA, they should not be affected by SCLB treatment and they should disappear completely after DNase I treatment. SCLB followed by DNase I reduced accessibility at clusters c1–c2 but not at c3; although these sites remain visible in our heatmaps, they are entirely absent from those of Yin et al. Considered on its own, the effect of SCLB followed by DNase I is consistent with their proposal that these sites arise from contaminating somatic cell DNA, lysed by SCLB and digested by DNase I. However, accessibility at clusters c1–c2 is partially restored by subsequent DTT treatment, which also increases accessibility at cluster c3 (Figures 7A and 7B). Such restoration is incompatible with the somatic-DNA model, since DNA that had been digested by DNase I and removed during the subsequent washes of sperm purification could not be regenerated by DTT. Representative changes at individual accessible sites are shown in Figure 7D.

To test the origin of the c1–c2 sites more directly, we reasoned that sites derived from a somatic genome should lack proteins specific to sperm chromatin. Using the same anchors, we examined the distribution of the sperm-specific histone variants TH2A and TH2B and of Protamine 1 from published ChIP-seq data ^10,44^. All three clusters, including c1–c2, were enriched for both sperm-specific histone variants and for Protamine 1, indicating that the c1–c2 sites originate from sperm chromatin rather than from somatic-cell DNA. These analyses also revealed a distinct chromatin organization at cluster c3 that may underlie its reduced accessibility to Tn5. The c1–c2 sites are flanked by nucleosomes and are themselves devoid of Protamine 1, which is instead enriched immediately outside the two flanking nucleosomes and abutting them. The c3 sites, by contrast, appear to be occupied by a single nucleosome that overlaps Protamine 1 (Figure 7E and Figure S7A).

The two classes of site also differ in their histone variants and modifications. Cluster c3 regions contain the canonical histones H3.1 and H3.2 but little H3.3, whereas the nucleosomes flanking clusters c1–c2 are enriched in H3.3 (Figure 7F). These H3.3 nucleosomes carry the active modifications H3K4me1, H3K4me3, H3K27ac, and H3K9ac, whereas the c3 sites carry the repressive modifications H3K27me3 and H3K9me3 (Figure 7F). Together, these results indicate that the c1–c2 sites do not derive from contaminating somatic chromatin. Rather, they lie in more accessible regions of sperm chromatin that are preferentially affected by DNase I once SCLB has permeabilized the sperm nucleus and allowed the enzyme to enter. The c3 sites lie in regions less accessible to Tn5; they can nonetheless be detected in untreated sperm at higher enzyme concentrations, or at low Tn5 concentrations once DTT treatment allows the sperm chromatin to escape the nucleus.

## DISCUSSION

Findings reported here address the effects of commonly used purification treatments during the isolation of mammalian sperm and have important consequences for the study of sperm chromatin. Yin et al. ^32^ have proposed that cauda sperm preparations are substantially contaminated by somatic cell chromatin, that this contamination dominates genome-wide profiles, and that valid sperm chromatin data can be obtained only after pretreatment with somatic cell lysis buffer, DNase I, and elevated DTT. If accepted, this argument would invalidate much of the published literature on mammalian sperm chromatin organization. Our results do not support these conclusions. Instead, we find that properly purified cauda sperm preparations carry no detectable somatic or cell-free DNA contamination, and that the proposed pretreatment is not a benign purification step but a sequence of treatments that progressively damages the sperm genome, allows chromatin to escape from the nucleus, and disrupts the architectural organization of the sperm genome. The differences between profiles obtained with and without this pretreatment therefore cannot be considered as the difference between contaminated and decontaminated sperm. We suggest that the observed discrepancies represent the difference between intact sperm chromatin and chromatin that has been fragmented and structurally reorganized by the protocol itself.

Contamination by somatic chromatin is a real concern for caput sperm, where extracellular DNA released from the epididymal epithelium is well documented and accounts for a substantial fraction of the methylation signal ^31^. However, sperm from the cauda epididymis, which is used for most chromatin analysis, has been shown to have very low levels of contamination from somatic cells ^31^. The level of contamination in sperm preparations may depend on the ability of the operator and on small changes of standard purification protocols. We have previously reported contaminations of less than one somatic cell per 1000 sperm, which cannot explain the observations reported by Yin et al., since a contaminant present at this level cannot dominate a genome-wide profile of ATAC-seq or Hi-C. Therefore, the use of harsh treatments involving permeabilization with detergents followed by treatment with DNase I are not justified when the sperm are devoid of somatic cell DNA impurities. What the pretreatment appears to do instead is to damage the sperm that it is meant to purify.

DNase I alone leaves sperm nuclei intact, consistent with the protamine-condensed genome being inaccessible to the nuclease in an intact cell. Somatic cell lysis buffer alone produces no visible nuclear change. But the two in sequence distort a fraction of sperm heads and, more importantly, generate TUNEL signal within virtually all sperm nuclei and a 2.5-fold rise in mean signal intensity. The morphologically abnormal population identified by Hoechst staining is only what is visible by fluorescence microscopy, with more widespread effects seen when using TUNEL. We interpret these results by suggesting that SCLB renders the sperm nucleus permeable, and DNase I then cleaves the sperm genome in situ. The 3% of sperm with overtly distorted heads is a lower bound on the damage, not a measure of it. DTT completes the process, discharging some of the chromatin out of the nucleus into an extended fibrillar web in essentially every cell. These effects extend to some of the proteins that establish sperm chromatin architecture, and the pattern of their displacement is itself informative. SCLB permeabilizes the plasma membrane and releases the midpiece pools of Protamine 1, SMC1, and H3K4me3. The presence of these proteins in the midpiece is unexpected and they may be left behind during the process of sperm nuclear condensation, in which case their loss is unlikely to bear on the structure and organization of chromatin. DTT, by contrast, releases protamine–protamine cross-links and discharges chromatin from the nuclear compartment, with Protamine 1 following the extruded DNA while CTCF, SMC1 and H3K4me3 remain largely associated with the residual nuclear space. This differential behavior matters for interpreting the Hi-C results of Yin et al., who reported that pretreated sperm lacks the contact domains seen in somatic cells and is instead disorganized. The simplest explanation is that chromatin that has been evicted from the nucleus cannot retain the spatial organization of an intact nucleus, regardless of whether the underlying protein-DNA contacts survive. The reported loss of higher-order structure is a consequence of the protocol, not a property of native sperm chromatin, which prior work has shown to be folded into loops and compartments resembling those of somatic cells. The use of DTT at concentrations that can also strip zinc-coordinated and redox-sensitive factors from DNA compounds the problem, introducing a second route by which architectural information could be lost.

The remaining argument of the need for treatment with SCLB-DNase I-DTT treatment is the premise that intact, protamine-packaged sperm chromatin is inaccessible to Tn5, such that any ATAC-seq signal from untreated sperm must originate from contaminating somatic cells. Our ATAC-see results refute this premise directly. Tn5 inserts fluorescent adapters throughout the nuclei of untreated cauda sperm, across the entire nuclear volume and in cells whose elongated morphology is unambiguously that of mature sperm and is readily distinguished from the round, larger nuclei of any somatic contaminant. No signal appears outside the sperm heads, arguing against a pool of extracellular somatic DNA. Sperm chromatin is therefore accessible to Tn5 without disulfide reduction, and ATAC-seq peaks obtained from untreated sperm reflect insertion into sperm chromatin rather than into contaminating material. The DTT step does not unlock an otherwise inaccessible substrate. Instead, it relocates the substrate, moving some of the chromatin into a disorganized, extranuclear form whose ATAC-see reactivity no longer corresponds to a native nuclear context.

Our analysis of the clustered ATAC-seq data reinforces the conclusion that accessible sites correspond to sperm chromatin rather than somatic cell contamination and exposes the internal inconsistency of the contamination model. The sites Yin et al. assign to contaminating somatic DNA (clusters C1–C2) carry the sperm-specific histone variants TH2A and TH2B and Protamine 1, marking them as sperm chromatin rather than somatic chromatin. Their accessibility is reduced by SCLB-plus-DNase I but partially restored by subsequent DTT. This restoration cannot be explained if digested somatic cell DNA had been digested by DNase I and removed in subsequent washes. The sites Yin et al. assign to sperm (cluster C3) prove to be present in untreated sperm as well, detectable at higher Tn5 concentrations and distinguished by a chromatin organization, repressive in character, that renders them less Tn5-accessible at the enzyme levels Yin et al. used. The disappearance of these sites in their results is consequently a sensitivity issue rather than a contamination signal. Both classes of site are features of sperm chromatin and neither requires a somatic explanation.

Taken together, these results overturn the methodological case made by Yin et al. and carry a constructive message for the field. The concern that has driven adoption of the SCLB-DNase I-DTT protocol, that untreated sperm profiles are dominated by somatic contamination, is not warranted for properly purified cauda sperm. Published ATAC-seq, MNase-seq, ChIP-seq, and Hi-C analyses of untreated cauda sperm do not require systematic re-examination. The appropriate methodological priority is a sperm purification process that preserves nuclear integrity rather than aggressive pretreatment that destroys it. Only chromatin that remains within an intact nucleus can report faithfully on the organization that the sperm genome carries to the egg and the paternal regulatory information this organization conveys to the next generation.

## RESOURCE AVAILABILITY

### Lead contact

Further information and requests for resources and reagents should be directed and will be fulfilled by the Lead Contact Victor Corces.

### Materials availability

This study did not generate new unique reagents.

### Data and Code Availability

- Raw ATAC-seq data are available from NCBI’s Gene Expression Omnibus (GEO) and are publicly available as of the date of publication. The accession number for all the datasets reported in this paper is GSE337896. Reviewers can access these data using token oncvuceethazdmb.
- This paper does not report original code.
- Any additional information required to reanalyze the data reported in this paper is available from the lead contact upon request.

## ACKNOWLEDGMENTS

We would like to thank Dr. Xingqi Chen from Uppsala University for a generous gift of Tn5 to perform ATAC-see reactions. This work was supported by U.S. Public Health Service Award R01 ES027859 from the National Institutes of Health to VGC. The content is solely the responsibility of the authors and does not necessarily represent the official views of the National Institutes of Health.

## AUTHOR CONTRIBUTIONS

KT planned and performed experiments, analyzed data, and wrote the manuscript. VGC planned experiments and contributed to writing the manuscript.

## STAR★Methods

### EXPERIMENTAL MODEL AND SUBJECT DETAILS

#### Ethics statement

Experiments presented in this study make use of sperm isolated from mice. All experiments were conducted according to the animal research guidelines from NIH and all protocols for animal usage were reviewed and approved by the Institutional Animal Care and Use Committee (IACUC) of Emory University.

#### Mice models

Mice were maintained and handled in accordance with the Institutional Animal Care and Use policies at Emory University. Mice were housed in standard cages on a 12: 12 h light:dark cycle and given ad lib access to food and water. Healthy 8–11 week-old CD1 male mice (Charles River Labs) not involved in previous procedures were used for sperm isolation. No genotyping was performed.

### METHOD DETAILS

#### Isolation of mouse sperm

Euthanasia was performed by cervical dislocation and the epididymis was removed. Mature sperm were collected from the dissected cauda epididymis of 8-11-week-old CD1 mice (Charles River Labs). After removal of blood vessels and fat in PBS, the cauda epididymis was washed with fresh PBS, placed in Donners medium in a cell culture plate, and punctured several times with a needle. The punctured cauda and 10 ml of sperm suspension, were then transferred to a tube and sperm were allowed to swim up for 1 hr at 37^0^C ^15^. The upper 6 ml of the sperm swim-up fraction was collected, pelleted by centrifugation, and washed twice with excess 1 x PBS. Purity of sperm was determined by examination under a microscope after Hoechst staining. Sperm purified using this procedure are referred to throughout this study as “untreated”. The same isolation process was followed for the caput epididymis.

#### Sperm treatments

Four different treatments were evaluated on cauda sperm, based on the methods of Yin et.al. ^32^. Since centrifugation conditions and washing steps were not fully described in the original manuscript, these procedures were adapted and performed in a manner consistent with the reported experimental workflow. Somatic Cell Lysis Buffer (SCLB) treatment: Untreated, purified sperm were centrifuged at 600 x g for 7 min, and the resulting pellet was resuspended in 1 ml of freshly prepared SCLB buffer (0.01% SDS, 0.005% Triton X-100) and incubated on ice for 10 min. The reaction was diluted with 1 ml PBS, followed by centrifugation at 700 x g for 9 min. The pellet was then washed once with PBS and centrifuged again at 700 x g for 9 min, after which the supernatant was removed. DNase I treatment: Purified sperm were centrifuged at 600 x g for 7 min, and the DNase I reaction mixture (Qiagen, Cat# 79254), prepared in 1 x PBS according to the manufacturer’s instructions, was added directly to the resulting pellet. Following gentle mixing by flicking the tube, samples were incubated at room temperature for 10 min. The reaction was stopped by the addition of excess PBS, followed by centrifugation at 700 x g for 9 min. The pellet was then washed with PBS and centrifuged again at 700 x g for 9 min, after which the supernatant was removed. SCLB + DNase I treatment: Following SCLB treatment as described above, the DNase I reaction mixture was added directly to the washed sperm pellet, and the DNase I treatment was subsequently performed as described above. SCLB + DNase I + DTT treatment: Following SCLB + DNase I treatment as described above, the washed sperm pellet was resuspended in 50 mM DTT prepared in PBS, gently mixed, and incubated for 1 h at room temperature. The reaction was quenched by adding excess volume of 100 mM N-ethylmaleimide (NEM). Samples were then centrifuged at 700 x g for 9 min, the pellet was washed with PBS and centrifuged again at 700 x g for 9 min, after which the supernatant was removed.

#### Agarose gel electrophoresis

Genomic DNA was isolated from untreated and treated sperm samples. Sperm were lysed in lysis buffer (0.1 M Tris-HCl pH8, 0.2 M NaCl, 5 mM EDTA, 0.4% SDS), supplemented with 0.2 mg/ml Proteinase K and 150 mM DTT, followed by incubation at 55°C for 2.5 h with shaking at 600 rpm. Genomic DNA was purified by phenol: chloroform: isoamyl alcohol extraction followed by ethanol precipitation. Purified DNA was resuspended in TE buffer and 150 ng of DNA per sample were resolved on 0.5% agarose gels. λ DNA-Mono Cut Mix (New England Biolabs, N30195) was used as size standards. Gel images were acquired using the Gel Doc EZ Imager (Bio-Rad) and analyzed with Image Lab v6 software (Bio-Rad).

#### In situ TUNEL

To assess DNA fragmentation, 10 μl of untreated or treated sperm suspended in PBS, were deposited onto Superfrost Plus slides (Avantor) and allowed to air-dry at room temperature. Samples were fixed in 4% paraformaldehyde in PBS pH 7.4 for 1 h at room temperature, followed by three washes in PBS for 5 min each. Sperm were then permeabilized with 0.1% sodium citrate and 0.1% Triton X-100 for 2 min at 4°C and washed twice in PBS for 5 min each. TUNEL staining was completed using the *In Situ* Cell Death Detection Kit, Fluorescein (Roche Diagnostics) according to the manufacturer’s instructions. Samples were counterstained with Hoechst, mounted with Fluoromount-G (SouthernBiotech), coverslipped and sealed with CoverGrip Coverslip Sealant (Biotium). The stained slides were imaged for subsequent analysis.

#### Immunofluorescence microscopy

PBS-washed untreated or treated sperm (10 μl) were deposited onto Superfrost Plus slides (Avantor) and allowed to air-dry at room temperature. Slides were rehydrated in PBS for 5 min, and sperm were treated with 10 mM DTT and 100 mM Tris-HCl pH8.0 for 1 h at room temperature. After washing with PBS for 10 min, sperm were fixed in 4% paraformaldehyde for 10 min at room temperature, followed by three washes in PBS 10 min each. Cells were permeabilized with 0.2% Triton-X in PBS for 10 min at room temperature and blocked with 10% normal goat serum (NGS; Thermo Fischer Scientific) for 1 h at room temperature. After blotting the excess serum, samples were incubated overnight at 4°C with primary antibody diluted in 10% NGS. The following day, slides were washed three times in PBS for 5 min each and incubated for 1 h at room temperature with the appropriate secondary antibody diluted in 10% NGS. After three washes in PBS 10 min each, nuclei were counterstained with Hoechst, slides were mounted with Fluoromout-G (SouthernBiotech) and sealed with CoverGrip Coverslip Sealant (Biotium). Widefield fluorescence images were acquired using a Leica DMi8 microscope, and confocal images were acquired using a Leica Stellaris 5 confocal microscope.Primary antibodies included rabbit polyclonal H3K4me3 (abcam) at 1:250 dilution, rabbit polyclonal SMC1 (Bethyl Laboratories) at 1:500, rabbit polyclonal CTCF (Millipore) at 1:400 and mouse monoclonal anti-Protamine 1 (Hup1N; Briar Patch) at 1:1000 dilution. Secondary antibodies included Alexa Fluor 488-conjugated goat anti-rabbit, Alexa Fluor 488-conjugated goat anti-mouse, and Alexa Fluor 647-conjugated goat anti-rabbit antibodies, all used at 1:1000 dilution.

#### Assay for Transposase-Accessible Chromatin with visualization (ATAC-see)

PBS-washed untreated or treated sperm (10 μl) were deposited onto Superfrost Plus slides (Avantor), allowed to air-dry at room temperature and rehydrated in PBS for 5 min. ATAC-see was performed previously described ^45^ with minor modifications to recapitulate the ATAC-seq reaction performed in suspension. Briefly, sperm were permeabilized on ice for 3 min in RSB lysis buffer (10 mM Tris-HCl pH 7.4, 10 mM NaCl, 3 mM MgCl₂) supplemented with 0.1% NP-40, 0.1% Tween-20, and 0.01% digitonin. Lysis was terminated by the addition of excess RSB supplemented with 0.1% Tween-20, followed by incubation on ice for 12 min. Cells were then incubated in 50 μl transposition reaction mixture containing 2.5 μl of 2 nM fluorescent Tn5-ATTO-488 transposase (kindly provided by Dr. Xingqi Chen) and 0.01% digitonin at 37°C for 35 min in the dark. Tagmentation was terminated by washing sperm three times for 15 min each at 55°C with ATAC-see Washing Buffer (0.01% SDS, 50 mM EDTA in PBS). Following a final wash in PBS, nuclei were counterstained with Hoechst, mounted, sealed and imaged as described above.

#### Assay for transposase-accessible chromatin using sequencing (ATAC-seq)

ATAC-seq was performed using the Omni-ATAC protocol ^46^. Washed pellets from untreated or treated sperm were gently resuspended in PBS. Following sperm counting, nuclei were isolated from 100,000 sperm by centrifugation at 600 x g for 8 min at 4°C, followed by incubation on ice for 3 min in RSB (see above) supplemented with 0.1% NP-40, 0.1% Tween-20, and 0.01% digitonin. Lysis was terminated by the addition of RSB containing 0.1% Tween-20. Nuclei were pelleted at 500 x g for 12 min and resuspended in the transposition reaction mixture containing 0.01% digitonin, followed by incubation at 37°C for 35 min in a thermomixer with shaking at 1,000 rpm. The reaction was terminated by the addition of genomic DNA lysis buffer (0.1 M Tris-HCl pH8, 0.2 M NaCl, 5 mM EDTA, and 0.4% SDS) and genomic DNA was purified as described above. Libraries were amplified using 2x KAPA SYBR Fast ABI Prism kit (Kapa Biosystems) and 2.5 µM indexed primers under the following PCR conditions: 72°C for 5 min; 98°C for 30 s; and 13 cycles at 98°C for 10 s, 63°C for 30 s, and 72°C for 1 min.

#### Analysis of ATAC-seq and ChIP-seq data

All ATAC-seq libraries were sequenced on an Illumina NovaSeq X Plus platform using 150-bp paired-end sequencing. Adapter sequences were trimmed using Trim Galore (https://www.bioinformatics.babraham.ac.uk/projects/trim_galore/?utm_source=chatgpt.com). Paired-end reads were aligned to the mouse mm10 reference genome using Bowtie2 with default parameters, except -X 2000 ^47^. PCR duplicates were removed using Picard MarkDuplicates (Broad Institute, GitHub Repository. Picard Toolkit. http://broadinstitute.github.io/picard/; Broad Institute. Accessed March 3, 2021). BAM files were subsequently filtered to retain properly paired reads with mapping quality ≥30 while excluding reads mapping to mitochondrial DNA, chromosome Y, unplaced contigs, and ENCODE mm10 blacklist regions ^48^. To account for the Tn5 insertion offset, aligned ATAC-seq reads were shifted as + strands by +4 bp and – strands by −5 bp ^49^. ATAC-seq fragments were then separated by size into 50-115 bp fragments, corresponding to transcription factor-bound regions, and 180-247 bp fragments, corresponding to mono-nucleosomes ^42^. Peaks were called from the sub-nucleosomal fractions using MACS2 with parameters:--nolambda --nomodel ^50,51^. Normalized bigWig tracks were generated from filtered BAM files using deepTools bamCoverage (--normalizeUsing RPKM, --binSize 10, --ignoreForNormalization chrY chrM, and --minMappingQuality 30) ^52^ and biological replicates were merged. Peak-centered accessibility heatmaps were generated from the union of MACS2 summit regions identified in the Untreated, SCLB+DNase I and SCLB + DNase I + DTT libraries. Summits were used as anchors, and sub-nucleosomal accessibility within ±2 kb of each summit was quantified, binned, and normalized using RPKM with deepTools computeMatrix. Heatmaps were generated using plotHeatmap, and regions were grouped by k-means clustering (*k* = 3) ^52^. Nucleosomal occupancy heatmaps were generated using the same anchor regions and identical cluster order defined by the sub-nucleosomal accessibility heatmaps. Mono-nucleosomal signal was quantified over the same genomic intervals using identical parameters. Average mono-nucleosomal occupancy profiles were generated using DANPOS2 ^53^.

#### Quantification and statistical analysis

The percentage of sperm exhibiting distorted Hoechst-stained morphology was calculated from 500 sperm counted per untreated and SCLB+DNase I conditions. Standard errors were estimated assuming a binomial distribution.

Agarose gel images were quantified using Image Lab v6 (Bio-Rad). Band intensity was measured within the software-defined band boundaries (fuchsia boxes), whereas total lane intensity was quantified across the entire lane (teal boxes), extending from the top of the well to the bottom of the gel using a fixed lane width of 6.79 mm. Band and total lane intensities were normalized to the band intensity of the untreated sample set to 1 to visualize changes in high-molecular-weight DNA relative to the total DNA signal present in each lane.

TUNEL images were acquired using a Leica DMi8 microscope, and mean fluorescence intensity was quantified in Fiji (Image J) for each biological replicate. Following confirmation of normality using the Shapiro–Wilk test, pairwise comparisons were performed using unpaired, two-tailed Student’s t tests with Bonferroni correction for multiple comparisons. Statistical significance is indicated as ns, not significant; **p < 0.01. Relative TUNEL fluorescence intensity was calculated by normalizing the mean fluorescence intensity of each treatment to that of the untreated group, which was set to 1. Bars represent mean ± SEM.

## SUPPLEMENTAL INFORMATION

**Figure S1.**
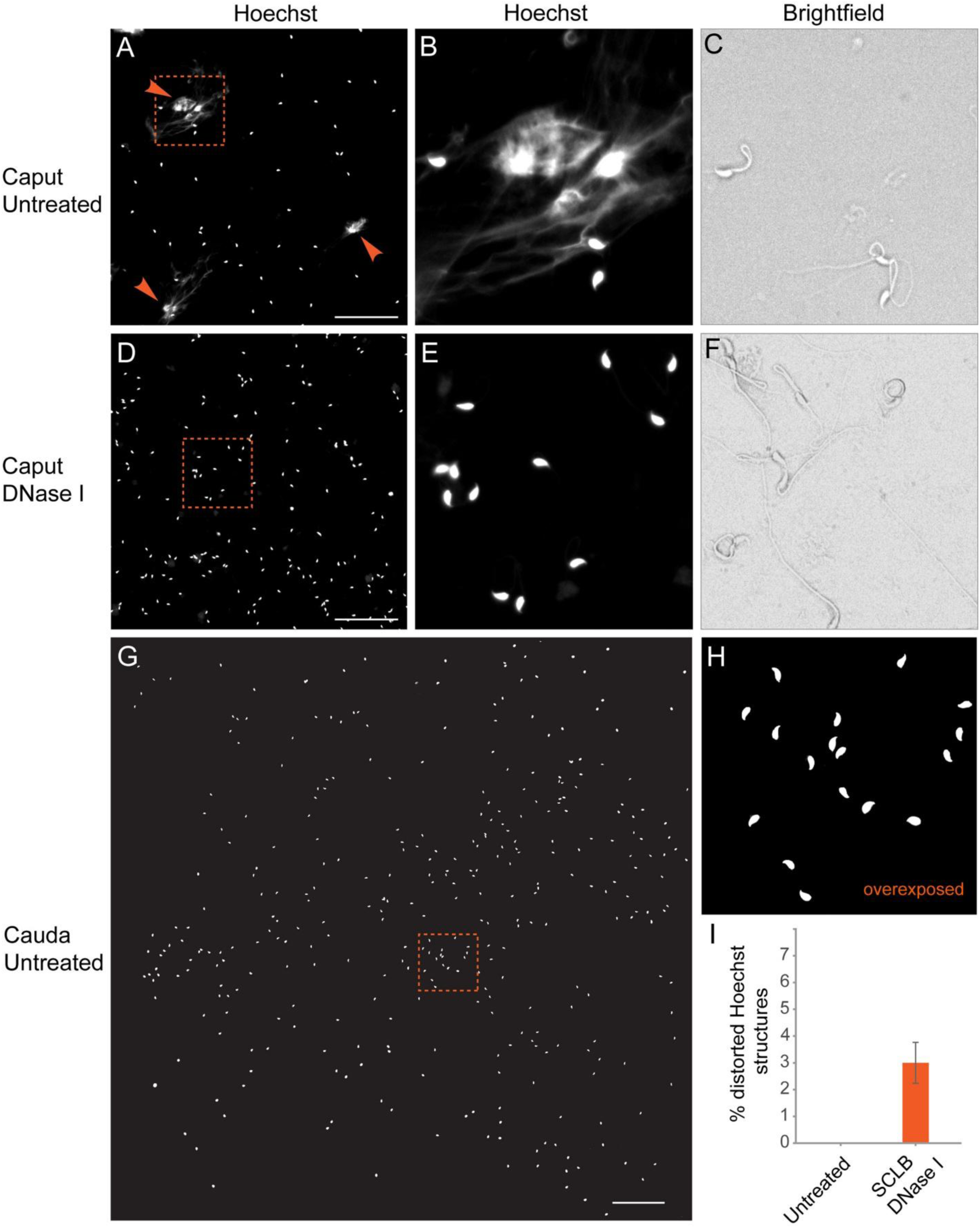
**Related to Figure 1** (A) Fluorescence microscopy of mouse sperm purified from the caput epididymis and stained with Hoechst. In addition to sperm heads, large masses of Hoechst-positive material can also be seen and are indicated by arrowheads. Scale bar corresponds to 150 μm. (B) Enlarged image of the area framed by the orange square in panel (A). (C) Brightfield microscopy of the area shown in panel (B). (D) Fluorescence microscopy of mouse sperm purified from the caput epididymis, treated with DNase I, and stained with Hoechst. The large masses of Hoechst-positive material cannot be seen after DNase I treatment. Scale bar corresponds to 150 μm. (E) Enlarged image of the area framed by the orange square in panel (D). (F) Brightfield microscopy of the area shown in panel (E). (G) Fluorescence microscopy of mouse sperm purified from the cauda epididymis and stained with Hoechst. Large masses of Hoechst-positive material or somatic cells cannot be seen in preparations of cauda sperm purified by the swim-up method. The image was automatically constructed by tiling multiple 20x-magnification images together. Scale bar corresponds to 200 μm. (H) Enlarged and over-exposed image of the area framed by the orange square in panel (G). (I) Quantification of Hoechst-positive distorted sperm heads observed in cauda sperm untreated or treated with SCLB followed by DNase I. All images are presented in grayscale to enhance structural detail. SCLB: Somatic Cell Lysis Buffer.

**Figure S2.**
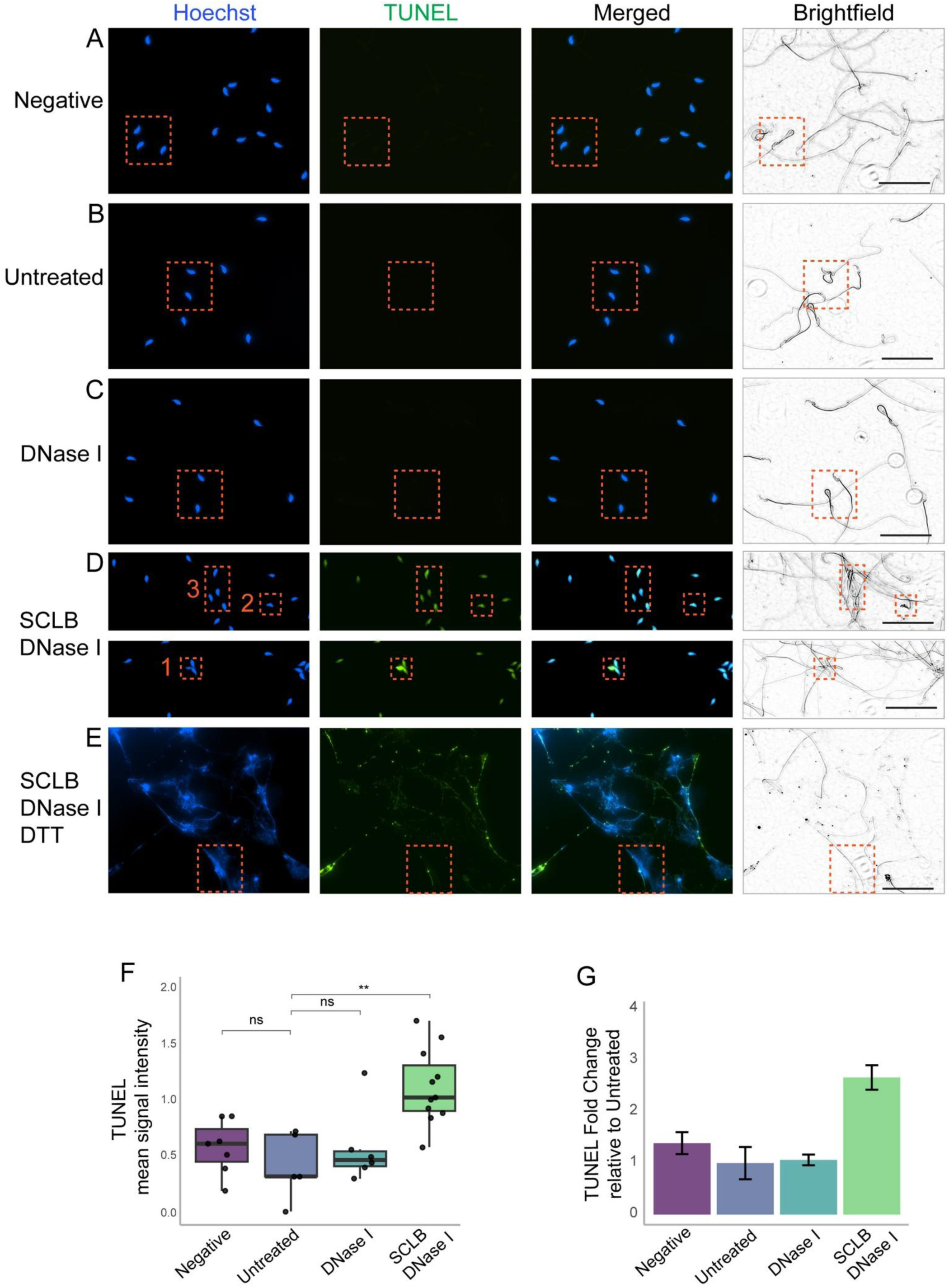
**Related to Figure 2** (A) Results from TUNEL assay and Hoechst staining of purified mouse sperm. Negative control lacking the TdT enzyme was included to assess background staining. Area framed by orange rectangle is shown in Figure 2D. Scale bar corresponds to 50 μm. (B) Results from TUNEL assay and Hoechst staining of purified mouse sperm. Untreated sperm. Area framed by orange rectangle is shown in Figure 2E. Scale bar corresponds to 50 μm. (C) Results from TUNEL assay and Hoechst staining of purified mouse sperm. Sperm treated with DNase I only. Area framed by orange rectangle is shown in Figure 2F. Scale bar corresponds to 50 μm. (D) Results from TUNEL assay and Hoechst staining of purified mouse sperm. Sperm treated with SCLB followed by DNase I. Areas framed by orange rectangles are shown in Figure 2G and labeled with the same numbers. Scale bar corresponds to 50 μm. (E) Results from TUNEL assay and Hoechst staining of purified mouse sperm. Sperm treated with SCLB followed by DNase I and then DTT. Area framed by orange rectangle is shown in Figure 2H. Scale bar corresponds to 50 μm. (F) Quantification of TUNEL mean signal intensity in sperm heads from sperm subjected to the treatments described at the bottom of the graph. Data are presented as mean ± SEM. (G) TUNEL signal intensity in sperm heads from sperm subjected to the treatments described at the bottom of the graph relative to untreated sperm. SCLB, Somatic Cell Lysis Buffer; DTT, Dithiothreitol; TUNEL, terminal deoxynucleotidyl transferase-mediated dUTP nick-end labeling; TdT, Terminal deoxynucleotidyl transferase; SEM, Standard Error of the Mean. **, p < 0.01; ns, not significant.

**Figure S3.**
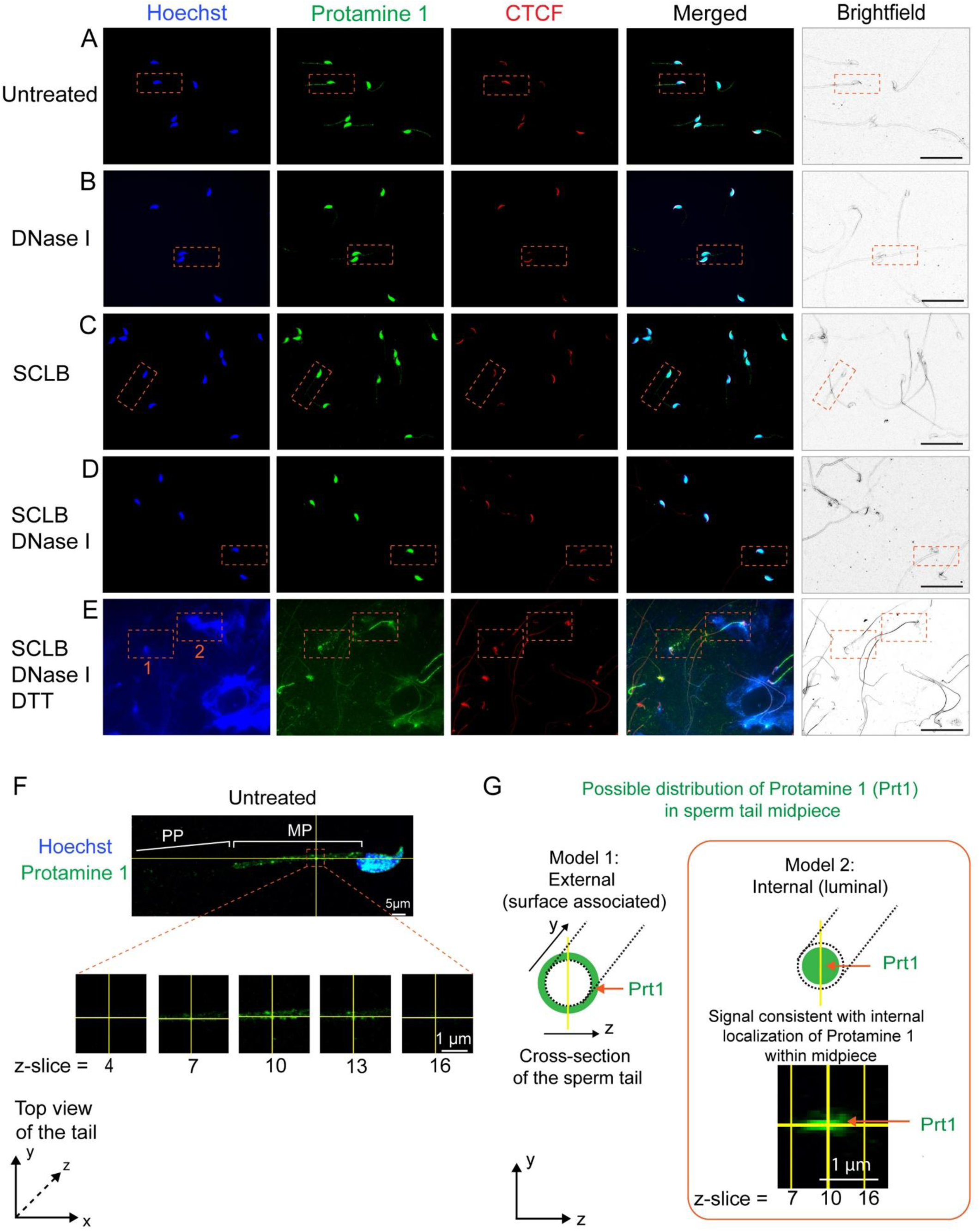
**Related to Figure 3** (A) Immunofluorescence and brightfield microscopy of purified untreated cauda sperm using antibodies to Protamine 1 and to CTCF. Area framed by an orange square is shown in Figure 3A. Scale bar corresponds to 50 μm. (B) Immunofluorescence microscopy of purified cauda sperm treated with DNase I using antibodies to Protamine 1 and to CTCF. Area framed by an orange square is shown in Figure 3B. Scale bar corresponds to 50 μm. (C) Immunofluorescence microscopy of purified cauda sperm treated with SCLB using antibodies to Protamine 1 and to CTCF. Area framed by an orange square is shown in Figure 3C. Scale bar corresponds to 50 μm. (D) Immunofluorescence microscopy of purified cauda sperm treated with SCLB followed by DNase I using antibodies to Protamine 1 and to CTCF. Area framed by an orange square is shown in Figure 3D. Scale bar corresponds to 50 μm. (E) Immunofluorescence microscopy of purified cauda sperm treated with SCLB followed by DNase I, and then DTT using antibodies to Protamine 1 and to CTCF. Areas framed by orange squares are shown in Figure 3E. Scale bar corresponds to 50 μm. (F) Serial confocal optical (z-) slices of the sperm midpiece stained with antibodies to Protamine 1. (G) 2D reconstruction compiled from consecutive confocal serial optical slices of Protamine 1 along the z-axis in the sperm tail midpiece, revealing internal luminal localization. SCLB, Somatic Cell Lysis Buffer; DTT, Dithiothreitol.

**Figure S4.**
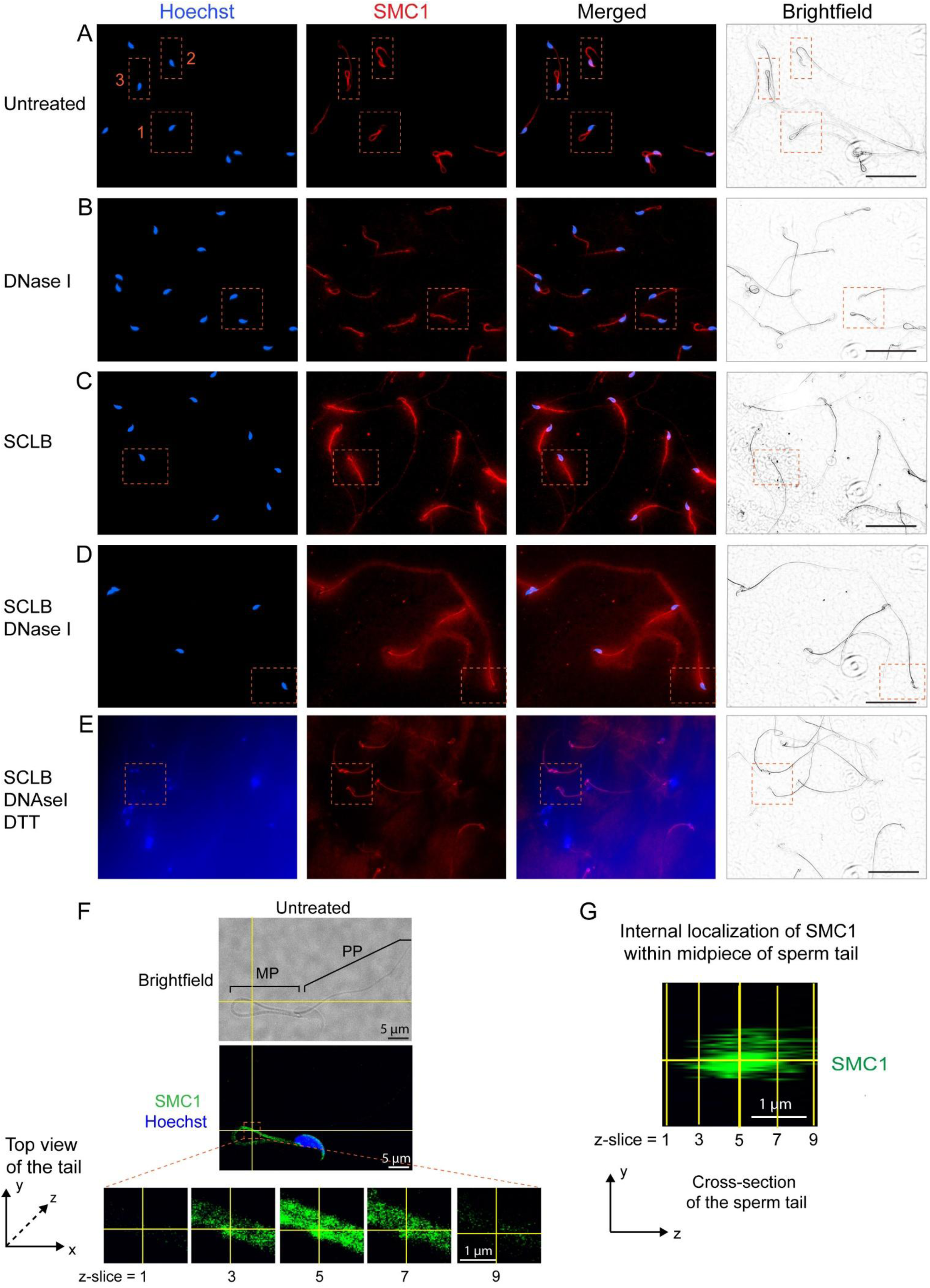
**Related to Figure 4** (A) Immunofluorescence microscopy of purified untreated cauda sperm using antibodies to SMC1. SMC1 shows nuclear staining, which ranges from low (panel 1) to high (panel 2) or intermediate (panel 3). SMC1 also shows very strong staining in the midpiece of the sperm tail. Areas framed by orange squares are shown in Figure 4A. Scale bar corresponds to 50 μm. (B) Immunofluorescence microscopy of purified cauda sperm treated with DNase I using antibodies to SMC1. Area framed by an orange square is shown in Figure 4B. Scale bar corresponds to 50 μm. (C) Immunofluorescence microscopy of purified cauda sperm treated with SCLB using antibodies to SMC1. Area framed by an orange square is shown in Figure 4C. Scale bar corresponds to 50 μm. (D) Immunofluorescence microscopy of purified cauda sperm treated with SCLB followed by DNase I using antibodies to SMC1. Area framed by an orange square is shown in Figure 4D. Scale bar corresponds to 50 μm. (E) Immunofluorescence microscopy of purified cauda sperm treated with SCLB followed by DNase I, and then DTT using antibodies to SMC1. Area framed by an orange square is shown in Figure 4E. Scale bar corresponds to 50 μm. (F) Serial confocal optical (z-) slices of the sperm midpiece stained with antibodies to SMC1. (G) 2D reconstruction compiled from consecutive confocal serial optical slices of SMC1 along the z-axis in the sperm tail midpiece, revealing internal luminal localization. SCLB, Somatic Cell Lysis Buffer; DTT, Dithiothreitol.

**Figure S5.**
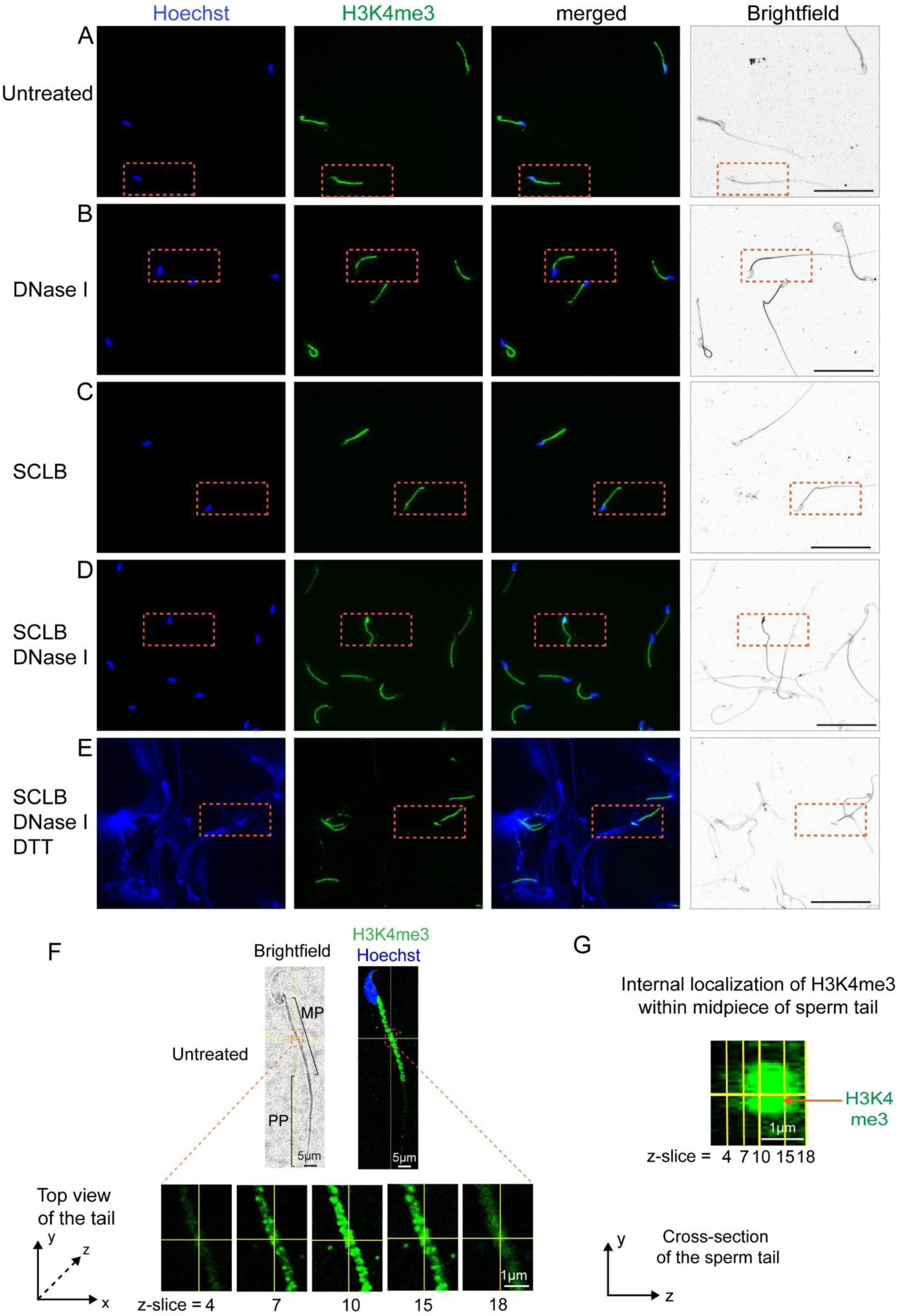
**Related to Figure 5** (A) Immunofluorescence microscopy of purified untreated cauda sperm using antibodies to H3K4me3. Area framed by an orange square is shown in Figure 5A. Scale bar corresponds to 20 μm. (B) Immunofluorescence microscopy of purified cauda sperm treated with DNase I using antibodies to H3K4me3. Area framed by an orange square is shown in Figure 5B. Scale bar corresponds to 20 μm. (C) Immunofluorescence microscopy of purified cauda sperm treated with SCLB using antibodies to H3K4me3. Area framed by an orange square is shown in Figure 5C. Scale bar corresponds to 20 μm. (D) Immunofluorescence microscopy of purified cauda sperm treated with SCLB followed by DNase I using antibodies to H3K4me3. Area framed by an orange square is shown in Figure 5D. Scale bar corresponds to 20 μm. (E) Immunofluorescence microscopy of purified cauda sperm treated with SCLB followed by DNase I, and then DTT using antibodies to H3K4me3. Area framed by an orange square is shown in Figure 5E. Scale bar corresponds to 20 μm. (F) Serial confocal optical (z-) slices of the sperm midpiece stained with antibodies to H3K4me3. (G) 2D reconstruction compiled from consecutive confocal serial optical slices of H3K4me3 along the z-axis in the sperm tail midpiece, revealing internal luminal localization. SCLB, Somatic Cell Lysis Buffer; DTT, Dithiothreitol.

**Figure S6.**
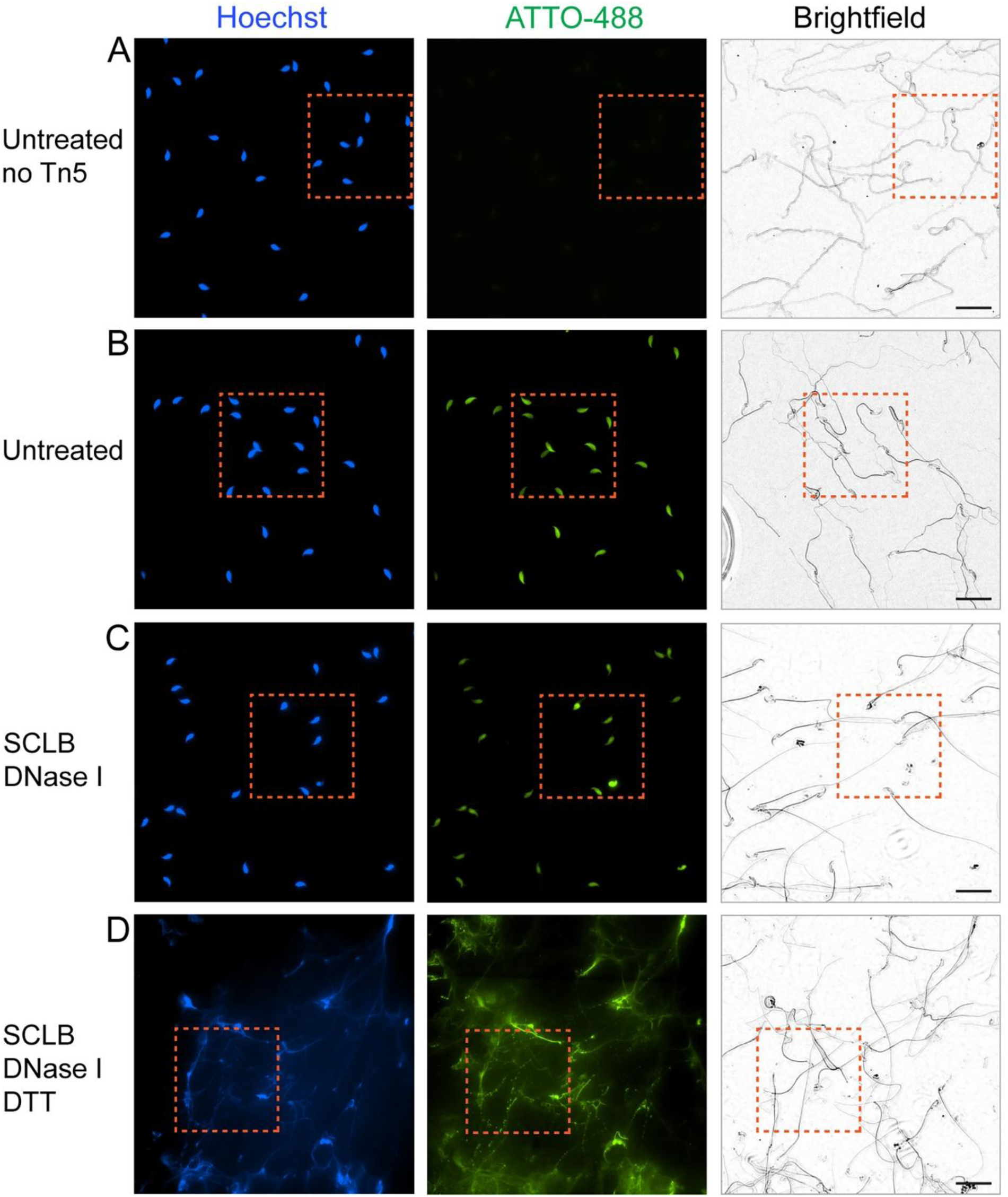
**Related to Figure 6** (A) Fluorescence microscopy of purified untreated cauda sperm subjected to the ATAC-see procedure with no Tn5 added. Area framed by an orange square is shown in Figure 6A. Scale bar corresponds to 25 μm. (B) Fluorescence microscopy of purified untreated cauda sperm subjected to the ATAC-see procedure. Area framed by an orange square is shown in Figure 6B. Scale bar corresponds to 25 μm. (C) Fluorescence microscopy of purified cauda sperm treated with SCLB followed by DNase I and subjected to the ATAC-see procedure. Area framed by an orange square is shown in Figure 6C. Scale bar corresponds to 25 μm. (D) Fluorescence microscopy of purified cauda sperm treated with SCLB followed by DNase I and then DTT. The sperm sample was then subjected to the ATAC-see procedure. Area framed by an orange square is shown in Figure 6D. Scale bar corresponds to 25 μm. SCLB, Somatic Cell Lysis Buffer; DTT, Dithiothreitol.

**Figure S7.**
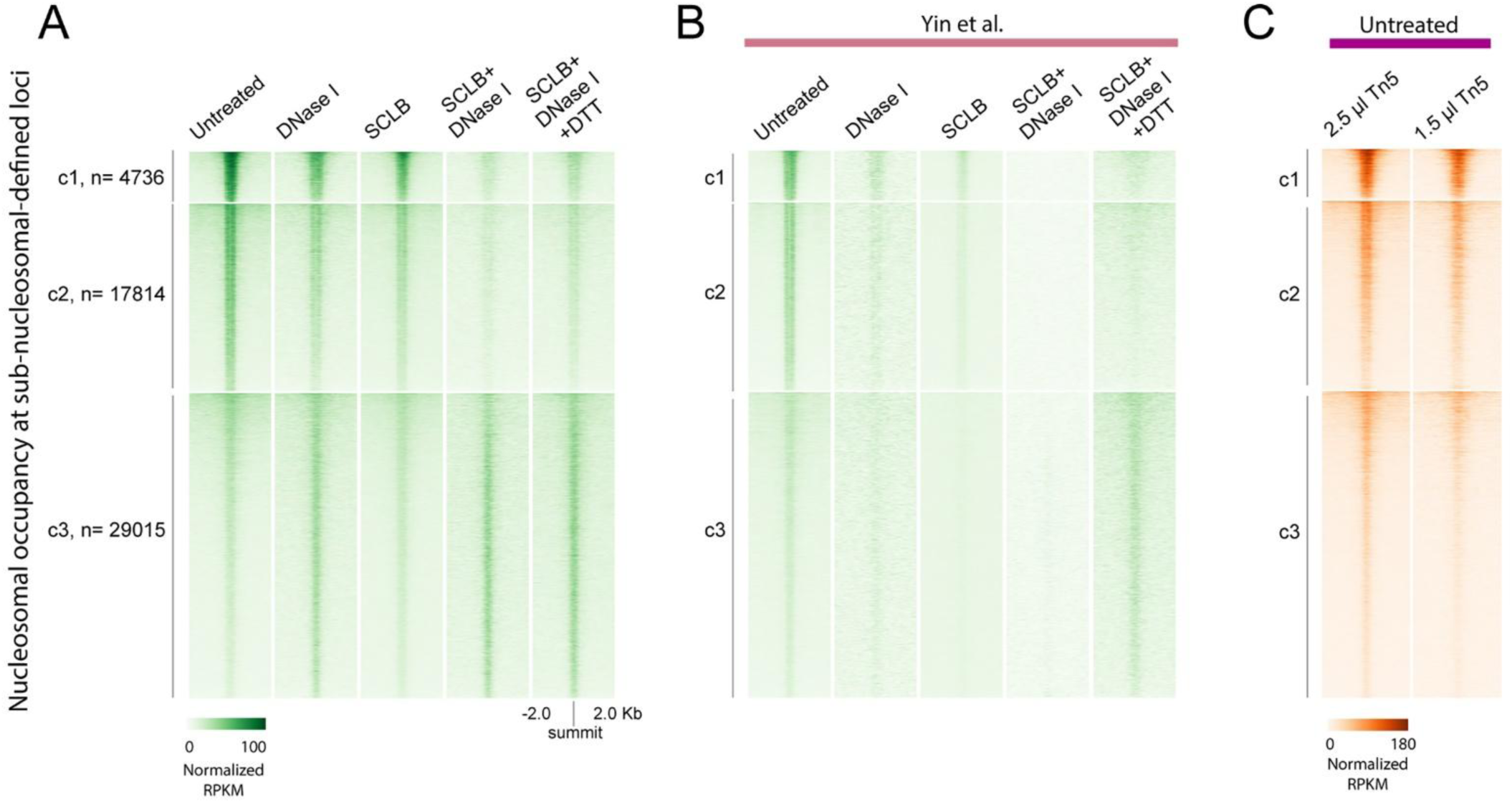
**Related to Figure 7** (A) Heatmaps of results from ATAC-seq experiments performed with purified cauda sperm. Sperm preparations were subjected to the treatments indicated above each of the heatmaps followed by the ATAC-seq procedure. Anchors are summits of peaks of sub-nucleosome size reads. Signal corresponds to mono nucleosome-size reads. (B) Heatmaps of results from ATAC-seq experiments performed with cauda sperm by Yin et al and downloaded from GEO accession GSE226685. Sperm preparations were subjected to the treatments indicated above each of the heatmaps followed by the ATAC-seq procedure as described by Yin et al. Data analysis was performed exactly as for the results described in panel (A). Anchors are summits of peaks of sub-nucleosome size reads. Signal corresponds to mono nucleosome-size reads. (C) Results of ATAC-seq experiments performed with purified untreated cauda sperm using 2.5 μl and 1.5 μl of Tn 5 transposase. Anchors are summits of peaks of sub-nucleosome size reads. Signal corresponds to mono nucleosome-size reads.

